# The oncogenic Runx3–Myc axis defines *p53*-deficient osteosarcomagenesis

**DOI:** 10.1101/2021.05.03.442316

**Authors:** Shohei Otani, Yuki Date, Tomoya Ueno, Tomoko Ito, Shuhei Kajikawa, Keisuke Omori, Ichiro Taniuchi, Masahiro Umeda, Toshihisa Komori, Junya Toguchida, Kosei Ito

## Abstract

Osteosarcoma (OS) in human patients is characterized by genetic alteration of *TP53*. Osteoprogenitor-specific *p53*-deleted mice (*OS* mice) have been widely used to study the process of osteosarcomagenesis. However, the molecular mechanisms responsible for the development of OS upon p53 inactivation remain largely unknown. In this study, we detected prominent RUNX3/Runx3 expression in human and mouse *p53*-deficient OS. Myc was aberrantly upregulated by Runx3 via mR1, a consensus Runx site in the *Myc* promoter, in a manner dependent on *p53* deficiency. Reduction of the Myc level by disruption of mR1 or Runx3 knockdown decreased the tumorigenicity of *p53-*deficient OS cells and effectively suppressed OS development in *OS* mice. Furthermore, Runx inhibitors exerted therapeutic effects on *OS* mice. Together, these results show that *p53* deficiency promotes osteosarcomagenesis in human and mouse by allowing Runx3 to induce oncogenic Myc expression.

## Introduction

*TP53* is the most frequently mutated gene in all types of human cancer, with mutations present in more than half of all tumors, and alteration of p53 that leads to loss of wild-type p53 activity is a master driver in most cancers (*1*–*3*). Accordingly, p53 is one of the most intensively studied tumor suppressor proteins. *p53-*null mice develop tumors at high penetrance, and *p53*-deficient mice crossed with mouse lines in which other cancer-related genes have been targeted have been widely used to elucidate the mechanisms of human cancer development. However, the diversified functions of p53 and the disparate consequences of its disruption prevent us from understanding the nature of *p53-*deficient carcinogenesis.

Osteosarcoma (OS) is the most common malignant bone tumor (*4*). Patients with germline mutations in *TP53* (Li-Fraumeni syndrome) have a high incidence of OS (*5, 6*), and *TP53* inactivation is often detected in sporadic OS (*4, 7*). In mice, restrictive deletion of *p53* in osteoprogenitor and mesenchymal stromal cells (MSCs) leads to development of OS with close histopathological resemblance to human OS, e.g., in the *Osterix* (*Osx*)*/Sp7*-Cre; *p53*^fl/fl^ mouse line, which is widely used as an animal model of human OS (*8, 9*). Thus, loss of p53 is a predominantly critical ‘solo-driver’ of osteosarcomagenesis in both human and mouse. Therefore, the scrutiny of *p53*-deficient osteosarcomagenesis should provide molecular insights into the universal mechanisms of tumorigenesis and malignancy caused or triggered by *p53* deficiency. Currently, however, little is known about the molecular events that result from loss of p53 and lead to OS development.

Dysregulation of transcription factors (TFs) plays pivotal roles in multiple cancers (*10*). Focusing on changes in the expression of genes encoding TFs, we compared the transcriptome of *p53*-deficient OS tissues to that of normal (wild-type) osteoblasts in human and mouse. Runx3, a member of the Runx family of genes, was the most upregulated TF in the absence of p53; c-Myc (Myc) and AP1 TFs, which have been attracted attention as oncogenes in OS (*4, 11, 12*), were also upregulated. Subsequent comprehensive genome-wide analyses revealed that Runx3 directly upregulates Myc in the *p53*-null context.

Our findings demonstrate that Runx3 functions as an oncogene to upregulate Myc via mR1, a genome element found in the *Myc* promoter; consistent with this, Runx inhibitors are efficacious against *p53*-deficient osteosarcomagenesis *in vivo*. Based on our findings, we propose that tumorigenesis driven by *p53* deficiency intrinsically requires the Runx3–Myc oncogenic axis.

## Results

### Runx3 is highly upregulated and oncogenic in *p53*-deficient OS in human and mouse

p53 inactivation is critical for osteosarcomagenesis in both human and mouse. Almost all OS patients (85 of 86) in the Therapeutically Applicable Research to Generate Effective Treatments (TARGET) cohort possessed *TP53* alterations: 84 had *TP53* genetic alterations, and one had *MDM2* amplification (Fig. 1A). *Osterix* (*Osx*)*/Sp7*-Cre; *p53*^fl/fl^ mice (herein *OS* mice) frequently developed OS as reported (*8, 9*) (fig. S1A). We compared the transcriptome of human or mouse OS tissues to that of normal osteoblasts or newborn calvaria, respectively (fig. S1B), focusing on changes in the expression of genes encoding 794 human TFs and 741 mouse orthologues. This analysis identified 47 TFs that were commonly up or downregulated more than 2-fold in human and mouse OS (Table S1). Seven of the top ten TFs (shown in blue; Fig. 1, A and B), when ranked by expression level, had orthologues in both species (fig. S1C). Among these seven TFs, *RUNX3* was associated with portended poor prognosis in human OS (fig. S1D). Moreover, *RUNX3* was over-expressed in almost every human and mouse OS; the highest degree of upregulation (log_2_FC) was 5.0 and 2.2 in human and mouse, respectively (Fig. 1, A and B). A similar, though less pronounced, trend was also observed for *MYC, JUNB*, and *FOS*. These observations inspired us to investigate the oncogenic roles of RUNX3/Runx3 in OS development.

**Fig. 1.**
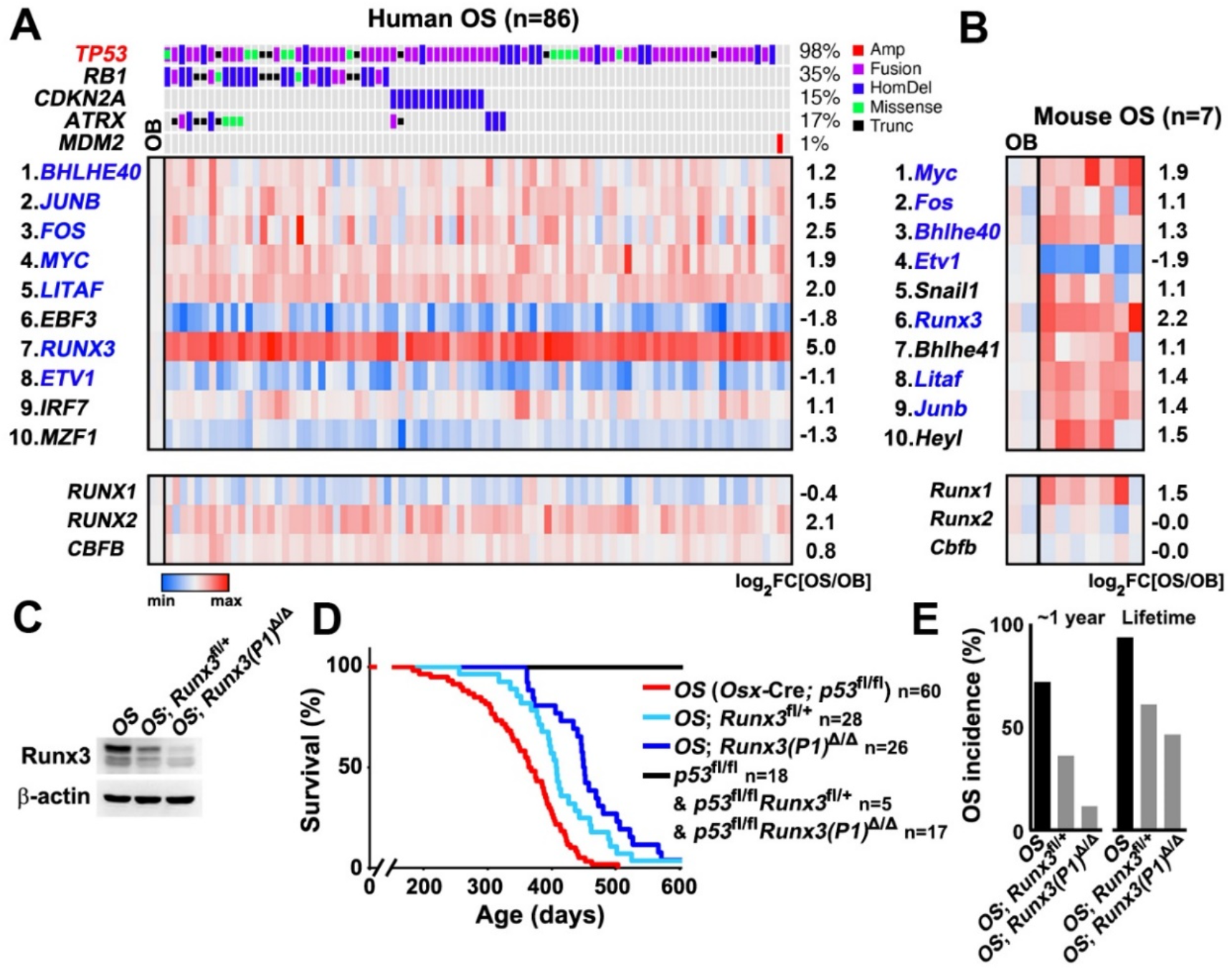
Runx3 is highly upregulated and oncogenic in *p53*-deficient OS. (**A**) Heatmap representing color-coded expression levels of the top 10 highest-expressed TFs and the RUNX family genes across 86 OS patients harboring genetic alterations in *TP53, RB1, CDKN2A, ATRX*, and *MDM2*. The ratio of gene expression level in OS vs. normal osteoblasts (OB) is shown as log_2_FC. (**B**) Heatmap of the top 10 highest-expressed TFs and the Runx family genes in OS in seven *OS* mice. Seven TFs common to both human and mouse are shown in blue (**A** and **B**). (**C**) Levels of the indicated proteins in BM-MSCs, as determined by western blotting. (**D**) Survival of *Runx3*^+*/*+^, *Runx3*^*fl/*+^, and *Runx3(P1)*^*Δ/Δ*^ mice in the *OS* background, alongside Cre-free controls. (**E**) Incidence of OS in the indicated genotypes within 1 year and throughout the lifespan.

RUNX3, a member of the RUNX family of genes, is a cancer-related TF(*13*). In contrast to the other members of the family, RUNX1 and RUNX2, and their subunit CBFβ, RUNX3 was markedly upregulated in both human (Fig. 1A) and mouse OS (Fig. 1B and fig. S2A). We isolated tumor cells from *OS* mice (mOS cells) and assessed their tumorigenic potential in immunocompromised mice (fig. S2B). mOS cell clones manifesting strong tumorigenicity were associated with aberrant Runx3 expression (fig. S2C) and recapitulated the histology of primary tumors (fig. S2D), in which Runx3 was specifically immunodetected in Runx2-positive osteogenic cells (fig. S2E). The positive correlation between tumorigenicity and RUNX3 expression was also observed in a series of human OS cells (fig. S3A). RUNX3/Runx3 knockdown decreased tumorigenicity in all human and mouse OS cells tested to a greater extent than knockdown of RUNX2/Runx2 (fig. S3, B to E). RUNX1/Runx1 expression had no significant effect upon osteosarcomagenesis (Fig. 1A, fig. S2, A and C, fig. S3A), and its knockdown did not affect the tumorigenicity of OS cells (fig S3B). Together, these results indicate that Runx3 is required for the tumorigenicity of *p53*-deficient OS cells.

We depleted *Runx3* from *OS* mice. Between the two independent promoters of *RUNX3/Runx3* (*14*), P1 and P2, *Runx3(P1)* was expressed several times more strongly than *Runx3(P2)* in mouse OS (fig S4A). *OS* mice with heterozygous *Runx3* deletion in osteoprogenitors (*OS*; *Runx3*^fl/+^) or systemic null mutation of *Runx3(P1)* (*OS*; *Runx3(P1)*^Δ/Δ^) (fig. S4, B to F) exhibited improvement in OS incidence and lifespan, mirroring the reduction of Runx3 expression in bone marrow (BM)-MSCs (Fig. 1, C to E). However, many *OS*; *Runx3*^fl/fl^ mice (*OS* mice lacking both *Runx3(P1)* and *Runx3(P2)* in osteoprogenitors) died before OS development (fig. S5, D and F), probably due to loss of the pro-proliferation function of Runx3 (*15*). By contrast, homozygous or heterozygous deletion of *Runx1* had little effect on lifespan and OS incidence in *OS* mice (fig. S5, A and B). *OS* mice with heterozygous deletion of *Runx2* (*OS*; *Runx2*^fl/+^), which is essential for osteoblast differentiation(*16*), died early before OS onset (fig. S5, C and F). Taken together, these results highlight the oncogenic roles of RUNX3/Runx3 in *p53*-deficient osteosarcomagenesis in human and mouse.

### Myc is a positive target of Runx3 in *p53*-deficient OS

To explore the target genes of Runx3 in *p53-*deficient OS cells, we investigated the genome-wide profiles of Runx3 and open and active chromatin using ChIP-seq of Runx3 and H3K27ac and assay for transposase-accessible chromatin using sequencing (ATAC-seq) (Fig. 2A) in a representative mOS cell line that exhibits Runx3-dependent tumorigenicity (fig. S3A). Genome-wide binding of Runx3 and open/active chromatin were highly concordant, with 1,624 genes strongly co-occupied by both Runx3 and accessible chromatin markers (or open/active chromatin), indicating that Runx3 acts as a general transcriptional activator in OS cells (Fig. 2A, fig. S6, A and B). In addition, we used microarrays to investigate changes in the transcriptome upon *Runx3(P1)* knockout in *p53*-null BM-MSCs; this analysis identified 1,552 potential targets upregulated by Runx3(P1) (Fig. 2B). Of the 95 genes shared between the 1,624 Runx3-occupied and 1,552 Runx3-upregulated genes (fig. S6C, table S2), *c-Myc* (*Myc*) was most strongly occupied by Runx3 and open/active chromatin within its regulatory region (Fig. 2A, table S2); moreover, this gene was among the factors most strongly associated with poor prognosis (fig. S6D). Thus, we identified Myc as the top candidate target gene upregulated by Runx3 in *p53*-deficient OS cells.

**Fig. 2.**
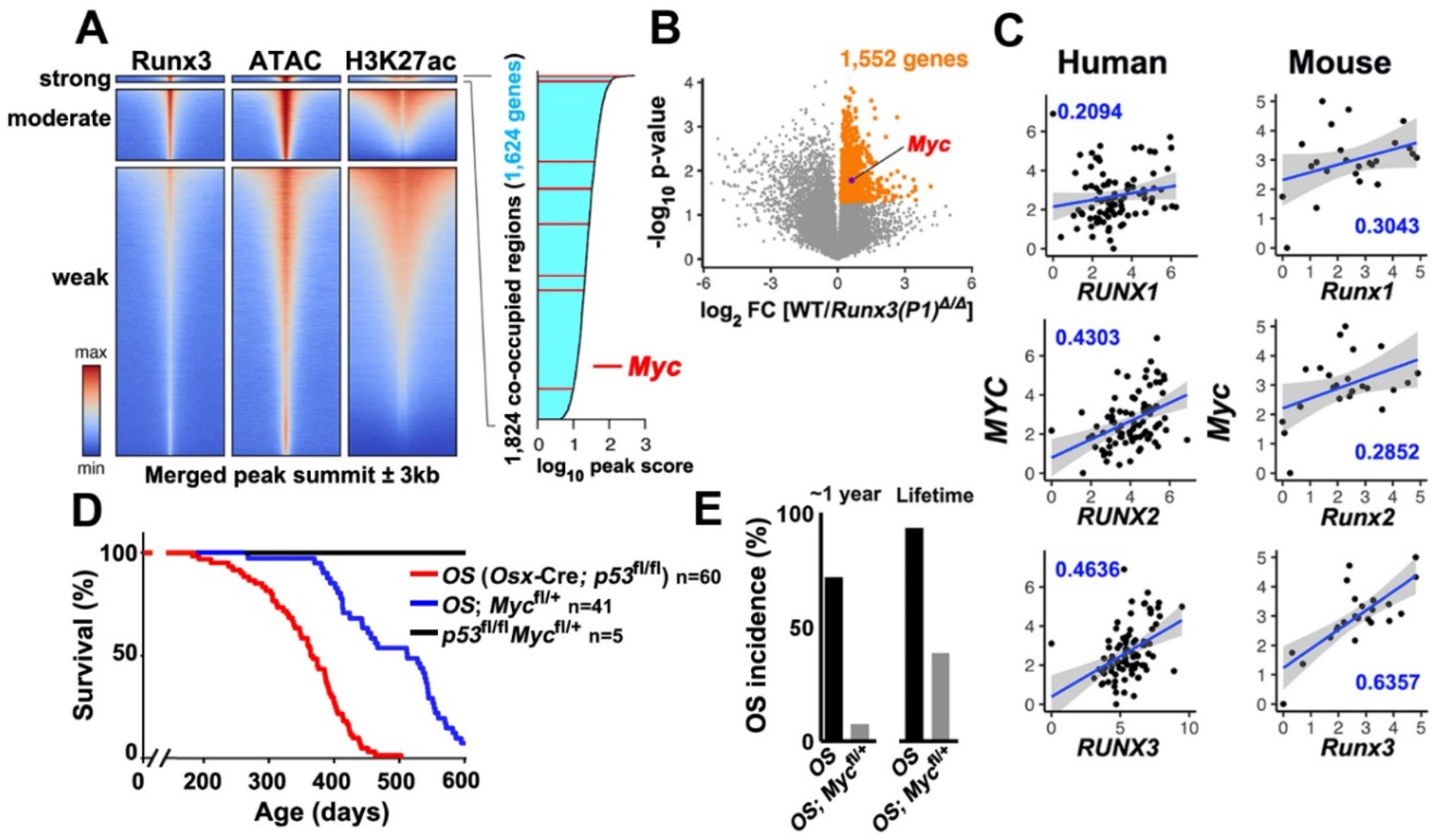
Myc is a positive target of Runx3 in *p53*-deficient OS. (**A**) Heatmaps showing genome-wide occupancy of Runx3, H3K27ac, and open chromatin (ATAC-seq). Regions (y-axis) were divided into three clusters, each ordered by merged signal intensity for all profiles. Myc regulatory regions (red) were most frequently observed among the 1,624 strongly co-occupied regions. (**B**) Volcano plot of the changes in gene expression upon Runx3(P1) deletion in BM-MSCs of *OS* mice. BM-MSCs from *OS* (WT; n=4) and *OS*; *Runx3(P1)*^Δ/Δ^ (*Runx3(P1)*^Δ/Δ^; n=5) mice were subjected to microarray analysis. Myc was one of the 1,552 genes (orange) significantly upregulated by Runx3(P1) in the absence of p53 (*p* < 0.05). (**C**) MYC/Myc expression levels plotted against RUNXs/Runxs expression levels in human (n=86, Fig. 1A) and mouse (n=24) OS. Spearman rank correlation coefficient is shown in each panel. (**D**) Survival of *Myc*^+/+^and *Myc*^fl/+^ mice in the *OS* background, alongside Cre-free controls. (**E**) OS incidence of each mouse line within 1 year and throughout the lifespan. The cohort of *OS* mice is identical to that in Fig. 1, D and E (**D** and **E**).

Myc, which is crucial for OS development (*17*), was markedly upregulated in human and mouse OS (Fig. 1, A and B). Among the RUNX family genes, the expression of *RUNX3/Runx3* was most strongly correlated with that of *MYC/Myc* in both human and mouse OS (Fig. 2C). Runx3 and Myc expression were highly correlated in terms of protein levels (fig. S7A) in mOS cells and localization in OS tissues (fig. S7, B and C). In human and mouse OS cells, knockdown of RUNX3/Runx3 led to down-regulation of MYC/Myc, whereas overexpression led to upregulation (fig. S7, D to F). Moreover, MYC/Myc was required for tumorigenicity (fig. S7, G and H). Importantly, heterozygous deletion of *Myc* greatly prolonged the lifespan and reduced OS incidence of *OS* mice (Fig. 2, D and E), as was the case for *Runx3* (Fig. 1, D and E). *OS* mice with homozygous deletion of *Myc* (*OS*; *Myc*^fl/fl^) died early before OS onset, as did *OS*; *Runx3*^fl/fl^ mice (fig. S5, D to F). Overall, the oncogenic effects of Myc and Runx3 were highly concordant in *p53*-deficient osteosarcomagenesis *in vivo*.

### The mR1 element is responsible for Myc upregulation by Runx3

To identify the elements within its 3-megabase (Mb) topologically associating domain (TAD) (*18*) (Fig. 3A) through which Runx3 upregulates *Myc*, we performed CRISPR-interference (CRISPRi) (*19*), in which chromatin is repressed by dCas9-KRAB in a targeted manner. For this analysis, we chose candidate elements with high levels of co-occupancy of Runx3 and open/active chromatin, as well as high conservation across mammals (Fig. 3A). The transcription start site (TSS) and leukemic enhancers N-Me and BDME of *Myc* (*20*) were also included as controls. Blockage of mR1, a consensus Runx site located ∼0.36 kb upstream of TSS, exhibited a significant reduction in Myc expression, comparable to the effect observed at the TSS in mOS cells (Fig. 3, B and C). None of the CRISPRi trials affected expression of *Pvt1*, a neighboring lncRNA that regulates *Myc* (*21*) (Fig. 3, A and C). All three RUNX consensus sites (MR1/mR1, MR2/mR2, and MR3/mR3) in the *MYC*/*Myc* promoter (fig. S8A) were bound by RUNX2/Runx2 and RUNX3/Runx3, but not RUNX1/Runx1 (fig. S8, B to F). Among them, only MR1/mR1 is in a region that is well conserved between human and mouse (fig. S8A). Consistent with this, blockage of only mR1 decreased the tumorigenicity and Myc expression level of mOS cells (Fig. 3, E and F), even though mR3 was predominantly bound by Runx3 (Fig. 3D and fig. S8, B and C).

**Fig. 3.**
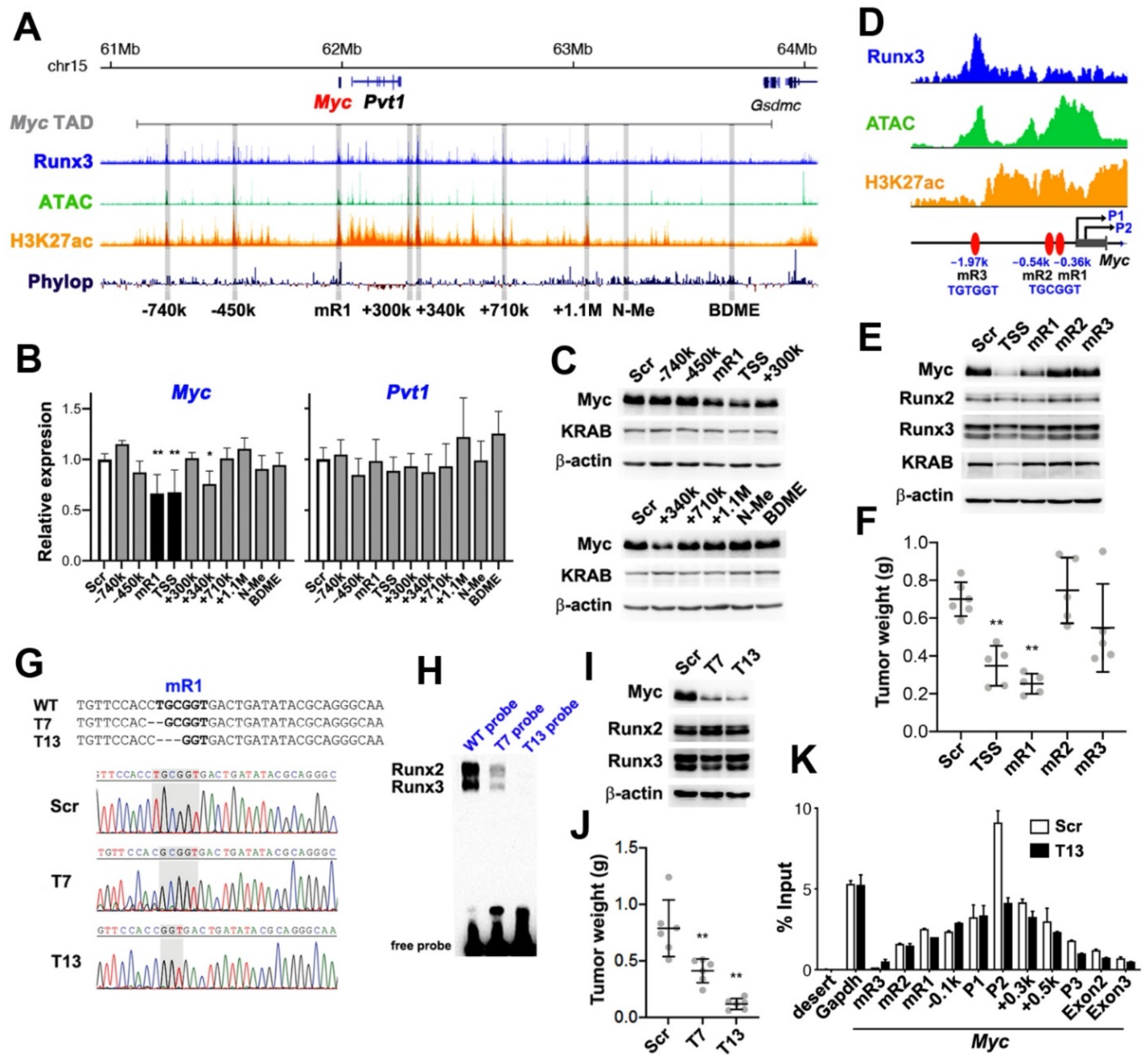
mR1 is a responsible element of Myc upregulation by Runx3. (**A**) Profiles of Runx3 and open/active chromatin (ATAC and H3K27ac) in mOS1-1 cells, together with the homology score (PhyloP), are aligned across the 3-Mb Myc TAD. Target regions of CRISPRi are shown in gray. (**B**) Relative expression of *Myc* and *Pvt1* in mOS1-1 cells in which the indicated regions were targeted by dCas9-KRAB. Scrambled (Scr) sgRNA served as a control. Data are means ± SD (n=3). ^**^*p* < 0.01; ^*^*p* < 0.05. (**C**) Levels of the indicated proteins in each CRISPRi mOS1-1 clone, as determined by western blotting. (**D**) Schematic representing mR1, mR2, and mR3 in the *Myc* promoter, with associated profiles for ChIP-seq (Runx3 and H3K27ac) and ATAC-seq. (**E**) Levels of the indicated proteins in CRISPRi mOS1-1 cells in which the indicated regions were targeted by dCas9-KRAB, as determined by western blotting. (**F**) Tumorigenicity of each CRISPRi mOS1-1 cells. (**G**) Sequence alignments of mOS1-1 clones with either 1 bp (T7) or 3 bp (T13) homozygous deletion in mR1, together with a non-targeted control (Scr). (**H**) EMSA performed using nuclear extract of mOS1-1 cells and labeled DNA probes with either intact (WT) or mutated (T7 or T13) mR1, corresponding to the sequences shown in (**G**). Specificity of probe-bound Runx2 or Runx3 is demonstrated in fig. S12E. (**I**) Levels of the indicated proteins in each genome-edited mOS1-1 clone, as determined by western blotting. (**J**) Tumorigenicity of each genome-edited mOS1-1 clone. (**K**) Occupancy of RNA polymerase II on the indicated positions of the *Myc* regulatory region, with positive (*Gapdh*) and negative (gene desert) controls, as revealed by ChIP in Scr and T13 mOS1-1 cells. Data are means ± SD (n=3).

To investigate the roles of mR1 with greater precision, we used genome editing to generate mutant mOS cell clones in which mR1 was homozygously disrupted (Fig. 3G). Depending on the degree of mR1 mutation that inhibited Runx2/3 binding, Myc expression and tumorigenicity of mOS cells were reduced (Fig. 3, G to J), as was also observed in another mOS cell line (fig. S9B) and human OS SJSA1 cells (fig. S9D). Deletion of a few bases neighboring mR1 or deletion of mR3 had little effect, although substitution of a single nucleotide within mR1 had a significant effect (fig. S9, A, C, E and F), confirming that mR1 was specifically required for oncogenicity. A mR1 mutation specifically inhibited entry of RNA polymerase II at the P2 promoter, from which the majority of *Myc* transcripts are derive (*22*), in mOS cells (Fig. 3K).

### mR1 and Runx3 are responsible for development of OS in *p53*-deficient mice

To evaluate the roles of mR1, mR2, and mR3 roles in mice, we replaced each of these sequences with a size-matched 6 bp *BglII* site using genome editing (fig. S10, E, H and I); mR1 was mutated using both homologous recombination and genome editing to ensure reproducibility (fig. S10, A to C, E). Mice harboring these mutations were crossed with *OS* mice (fig. S10D). Homozygous disruption of mR1, but not of mR2 or mR3, abolished Myc upregulation in BM-MSCs of *OS* mice (Fig. 4A). Regardless of how the mice were generated, homozygous disruption of mR1 (*OS*; *mR1*^m/m^) improved the lifespan and OS incidence of *OS* mice, whereas mutation of mR2 (*OS*; *mR2*^m/m^) or mR3 (*OS*; *mR3*^m/m^) did not have such a tumor-suppressive effect (Fig. 4, B and C, fig. S10, F, G, J, and K). In fact, *OS*; *mR1*^m/m^ mice were nearly identical to *OS*; *Runx3*^fl/+^ mice in terms of survival and OS incidence (Fig. 4, D and E). Together, these results clearly show that in a *p53*-deficient setting, Runx3 upregulates Myc via mR1 to promote OS development.

**Fig. 4.**
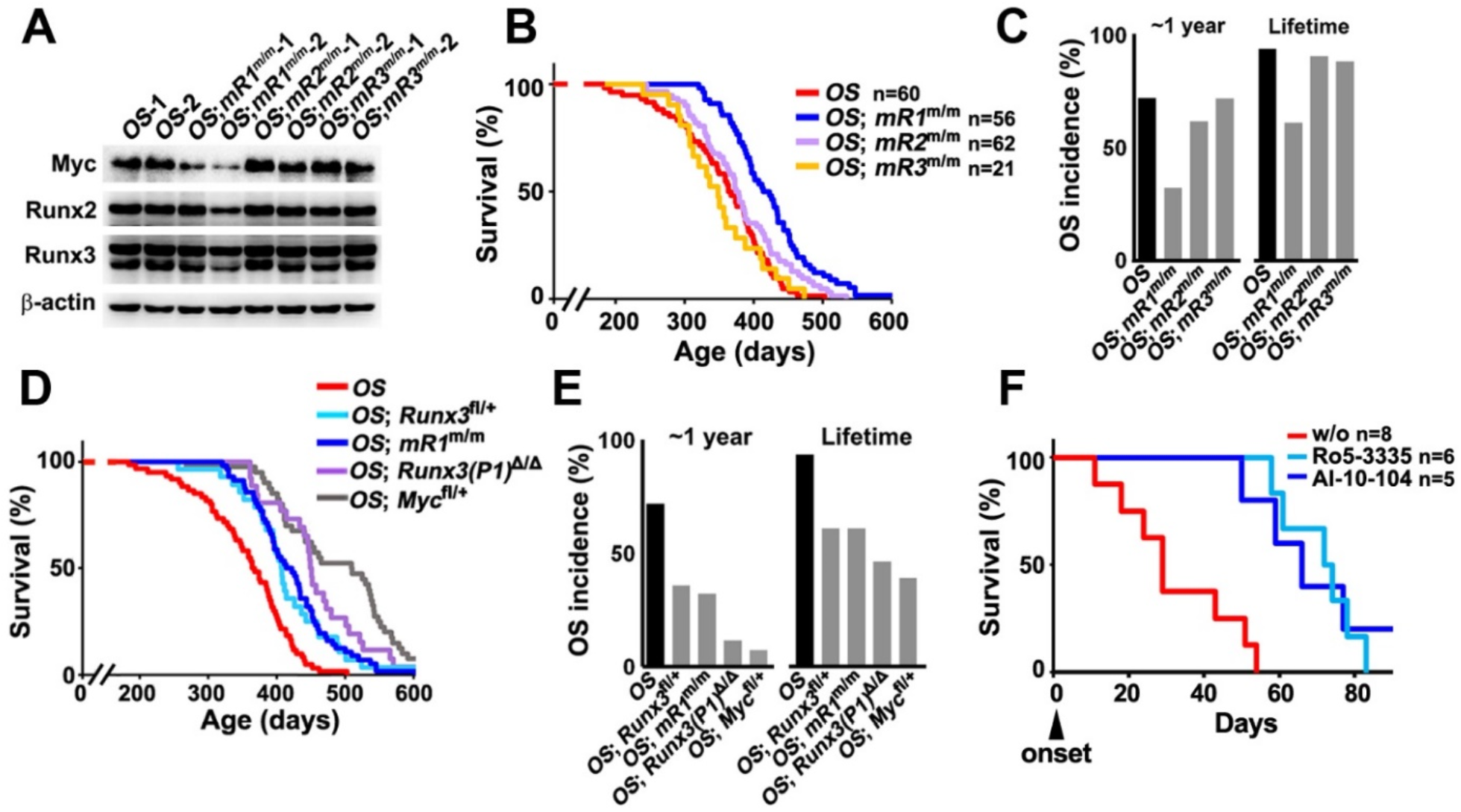
mR1 and Runx3 are responsible for development of OS in *p53*-deficient mice. (**A**) Levels of the indicated proteins in BM-MSCs from two individuals of each mouse line, as determined by western blotting. (**B**) Survival of the indicated genotypes. (**C**) Incidence of OS in the indicated genotypes within 1 year and throughout the lifespan. The results of *OS* mice are identical to those in Fig. 1, D or E, and those of *OS*; *mR1*^m/m^ and *OS*; *mR2*^m/m^ mice are from two independent lines, each shown in fig. S10, F, G, J, and K (**B** and **C**). (**D**) Comparison of survival of *OS* mouse lines shown in Fig. 1D, 2D, and (**B**): *OS, OS*; *Runx3*^fl/+^, *OS*; *mR1*^m/m^, *OS*; *Runx3(P1)*^Δ/Δ^, and *OS*; *Myc*^fl/+^. (**E**) Comparison of OS incidence in *OS* mouse lines shown in (**D**) within 1 year and throughout the lifespan. (**F**) Survival of *OS* mice treated with or without (w/o) Ro5-3335 or AI-10-104 after the onset of OS.

For further confirmation of the Runx3 oncogenicity, *OS* mice were treated with the Runx inhibitors, Ro5-3335 (*23*) and AI-01-104 (*24*), which inhibit the interaction between Runx and Cbfβ, thereby inhibiting Runx transactivation. Administration of either of these compounds effectively prolonged the lifespan of *OS* mice after the onset of OS (Fig. 4F).

### Induction of Myc by Runx3 is dependent on *p53* deficiency

In the absence of p53, disruption of either mR1 or Runx3(P1) prevented upregulation of Myc in BM-MSCs (Fig. 4A and Fig. 5B). On the other hand, in the presence of p53, neither disruption of mR1 nor disruption of Runx3(P1) affected Myc expression, which was expressed at a low basal level in BM-MSCs (Fig. 5, A and B). Interestingly, restoration of p53 significantly decreased Myc upregulation in *p53*-negative BM-MSCs, but did not reduce expression below the basal level of expression that was retained in the absence of Runx3 (Fig. 5C). Likewise, Myc was efficiently downregulated by p53 induction in mOS cells, but not in *Runx3*-negative mOS cells derived from OS that occasionally developed in *OS; Runx3*^fl/fl^ mice (fig. S11, A to C). Therefore, Runx3 upregulates Myc via mR1 only in the *p53*-negative context.

**Fig. 5.**
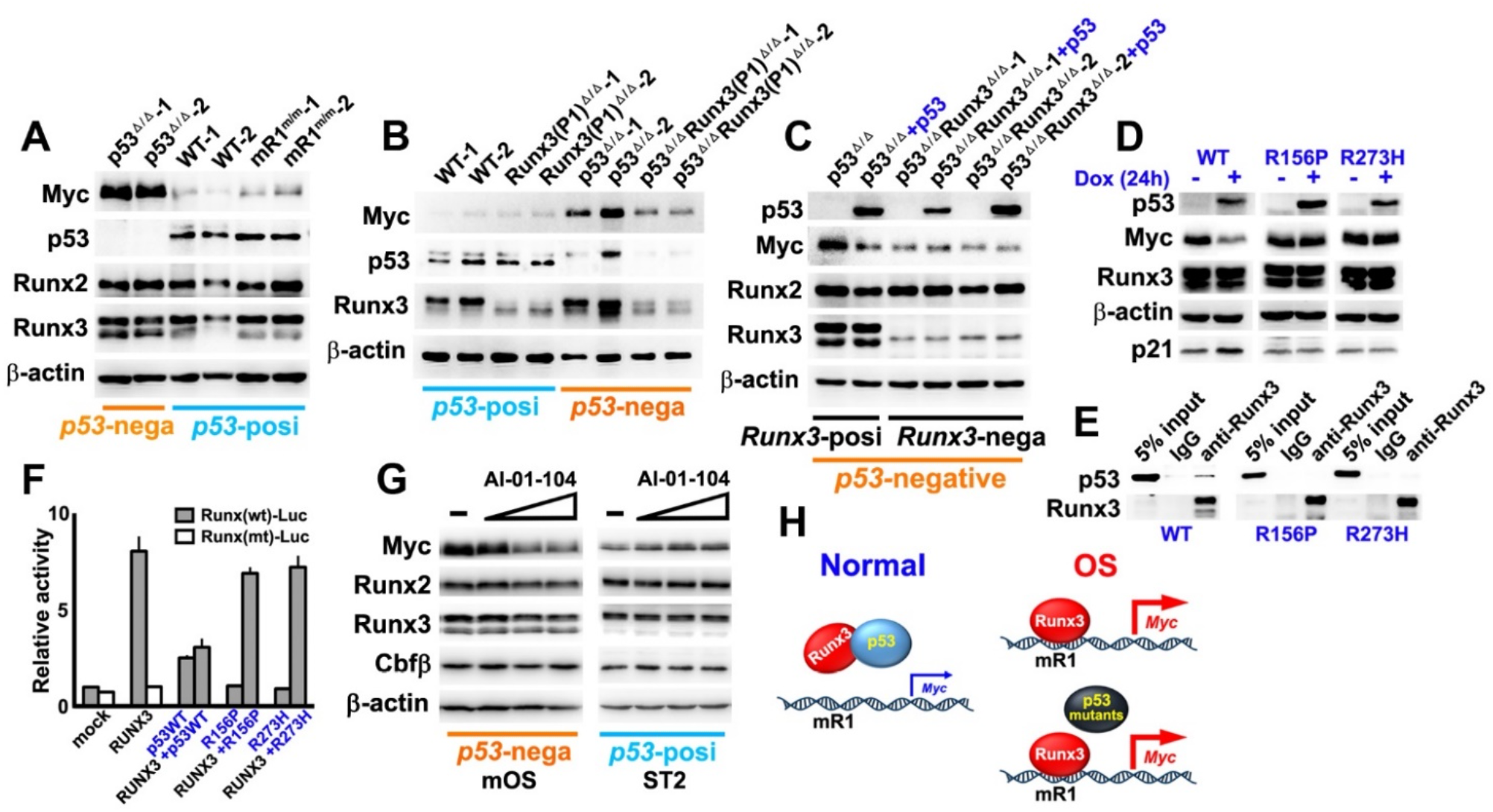
Myc induction by Runx3 is dependent on *p53*-deficiency. (**A**) Levels of the indicated proteins in BM-MSCs from two individuals of each mouse line, *OS* (p53^Δ/Δ^), wild-type (WT), or *mR1*^m/m^ (mR1^m/m^), as determined by western blotting. (**B**) Levels of the indicated proteins in BM-MSCs from two individuals of each mouse line [WT, *Runx3(P1)*^Δ/Δ^ (Runx3(P1)^Δ/Δ^), p53^Δ/Δ^, or *OS*;*Runx3(P1)*^Δ/Δ^ (p53^Δ/Δ^Runx3(P1)^Δ/Δ^) mice], as determined by western blotting. (**C**) Levels of the indicated proteins in p53^Δ/Δ^ or p53^Δ/Δ^Runx3(P1)^Δ/Δ^ BM-MSCs with or without exogenously restored p53, as determined by western blotting. (**D**) Levels of the indicated proteins in mOS1-1 cells expressing either WT or mutant p53 (R156P, R273H). Expression of p21 confirmed WT p53 function. (**E**) Co-immunoprecipitation of endogenous Runx3 and exogenous WT, R156P, or R273H p53 in mOS1-1 cells. Levels of the indicated proteins in immunoprecipitates obtained using anti-Runx3 antibody were determined by western blotting. (**F**) Effect of WT, R156P, or R273H p53 on RUNX3 transcriptional activity in *p53*- and *RUNX3*-negative G292 cells (n=3). (**G**) Efficacy of AI-10-104 (0, 0.2, 0.6, or 1.8 μM for 24 hours) on Myc expression in *p53*-negative mOS2-2 cells and *p53*-positive ST2 cells. (**H**) In normal cells, p53 attenuates Runx3 transactivation, whereas in OS, Runx3 aberrantly upregulates Myc in the absence of p53 or in the presence of mutant p53.

In human OS cells, exogenous RUNX3 induced MYC in *p53*-negative G292 cells but not in *p53*-positive U2OS cells (fig. S11D). Notably, two p53 mutants expressed in HOS/MNNG-HOS/143B (R156P) (*25*) and NOS-1 (R273H) cell lines (fig. S11E) failed to repress Myc (Fig. 5D and fig. S11F). Importantly, although Runx3 directly interacted with wild-type p53, as previously reported (*26*), both p53 mutants lost their interaction with Runx3 (Fig. 5E) and the ability to suppress Runx3 transcriptional activity (Fig. 5F). Consistent with this, MNNG-HOS and NOS-1 cells exhibited RUNX3-dependent MYC regulation (fig. S7F). These results suggest that p53 prevents Myc upregulation by physically inhibiting Runx3.

Next, we performed EMSA to determine whether p53 affects Runx3 ability to bind mR1. In these experiments, we used *p53*-negative mOS cells and other *p53*-positive murine cell lines: ST2 (a BM-MSC line) and 3T3-E1 (an osteoblast progenitor line). To quantitatively compare the amounts of endogenous Runx proteins, we utilized a pan-Runx monoclonal antibody that evenly reacted with all Runx proteins by recognizing the conserved C-terminal VWRPY motif (fig. S12, A to C). In ST2 and 3T3-E1 cells, both of which were *p53*-positive, the amount of mR1-bound Runx3 was smaller than in *p53*-negative mOS cells, and inversely proportional to the amount of endogenous p53 (fig. S12, C to E). This observation implies that p53 inhibits Runx3 DNA binding. The amount of mR1-bound Runx2, on the other hand, was constant and less strongly affected by the presence of p53 (fig. S12, C to E). In fact, the amount of Runx3 bound to mR1 was reduced by either addition of p53 (fig. S12F) or induction of p53 in mOS cells (fig. S12G), whereas the amount of bound Runx2 was unaffected. Consistent with these results, p53 exhibited a stronger interaction with Runx3 than with Runx2, both exogenously and endogenously (fig. S12, H and I), and more effectively attenuated the transcriptional activity of Runx3 than that of Runx2 (fig. S12J).

RUNX2 reportedly promotes development of OS (*13, 27, 28*). To assess the function of this protein, we examined *Runx3*-negative mOS cells and observed both Runx2- and mR1-dependent tumorigenicity. In these cells, the level of Myc correlated well with that of Runx2 (fig. S13A), and knockdown of Runx2 decreased both Myc expression and tumorigenicity (fig. S13B). In this *Runx3*-negative context, Runx2 bound to mR1 (fig. S13, C to E), which was responsible for Myc upregulation and tumorigenic potential in these cells (fig. S13, F and G), whereas Runx1 exhibited neither correlation with Myc expression nor occupancy of mR1 (fig. S13, A, C to E). Thus, Runx2 compensated for Runx3 in osteosarcomagenesis, and in OS cells with weaker Runx3 oncogenicity or more normal p53 activity, the pro-tumorigenic activity of Runx2 is more pronounced.

The Runx inhibitor, AI-01-104 decreased Runx3-enhanced Myc expression in *p53*-negative mOS cells, but did not affect the basal physiological level of Myc in *p53*-positive ST2 cells (Fig. 5G). The ability of Runx inhibitors to repress Myc was specific to the *p53*-deficient context, clearly demonstrating that induction of Myc by Runx3 is dependent on *p53* deficiency (Fig. 5H).

## Discussion

*p53-*deficient osteosarcomagenesis was inhibited by reduction of Myc or Runx3, disruption of mR1, or administration of Runx inhibitors, demonstrating that a key feature of *p53*-deficient osteosarcomagenesis is the aberrant upregulation of Myc by Runx3 via mR1. In this study, we did not address the mechanisms by which RUNX3/Runx3 was upregulated, however, it is not unreasonable to assume that oncogenic signaling pathways in the *p53*-deficient tumor microenvironment play a role. For instance, TGF-β signaling strongly induces oncogenic Runx3 upregulation in *p53*-deficient pancreatic ductal adenocarcinoma (*29*), in which Myc functions as a critical oncogene (*30*).

The tumor-suppressive function of RUNX3 initially attracted attention based on observations of gastric phenotypes in *Runx3-*deficient mice and the causal relationship between *RUNX3* silencing and the genesis of human gastric cancer (*31*). In a diverse range of human cancers, including gastric, colorectal, lung, pancreas, breast, liver, and prostate cancers, as well as leukemia and neuroblastoma, RUNX3 inactivation occurs mainly due to hypermethylation of the promoter or protein mislocalization (*13, 32*). More recently, on the other hand, RUNX3 upregulation has been also observed in various cases of human malignant tumors, suggesting its oncogenic roles (*13, 33*). Over the past two decades, research on RUNX3 further revealed its tumor-suppressive or oncogenic functions, bringing sharper focus on a fundamental question; how are the dualistic roles of RUNX3 determined by cellular context? Given the potential medical value of targeting RUNX transcriptional activities (*23, 34, 35*), the demand to answer this question is growing. The results presented indicate that p53 status is a contextual determinant of the dual roles of RUNX3 (*36*). p53 inactivation is the crucial event responsible for Runx3 oncogenicity leading to development of OS. In fact, in *p53*-positive U2OS cells (fig. S11D), elevated levels of RUNX3 induced p21 expression (data not shown), highlighting the fact that RUNX3 can play a tumor-suppressive role by promoting transactivation of wild-type p53, as previously reported (*26*).

As with loss of p53, MYC activation has been observed in more than half of all cancers; Myc directly contributes to malignant transformation through its pathogenic roles in tumor initiation, progression, and maintenance (*37*). *Myc* is critically regulated by tissue-specific regulatory regions, underscoring the fundamental importance of cancer-specific enhancers/superenhancers, and making *Myc* gene the best example thus far of long-range regulation. However, the fundamental mechanisms driving enhancer–promoter transactivation remain unclear (*38*), probably because the available evidence regarding TFs responsible for the transactivation remains inadequate. Under these circumstances, identification of mR1, a promoter element that is essential for aberrant upregulation of *Myc*, can provide deeper insight into the enhancer–promoter regulation of *Myc*. Given that Runx3 is a general and genuine transcriptional activator of Myc in the absence of p53, and that Runx1 is involved in *Myc* regulation via superenhancers (*20*), Runx3 may function as a crucial modulator to activate the superenhancer/core promoter of *Myc* in concert with other transcription factors such as Smads (*29*) and AP1, both of which interact with Runx proteins (*39*). AP1 TFs are prominently upregulated in human and mouse OS (Fig. 1, A and B), and, interestingly, consensus motifs of AP1 and Runx are co-enriched genome-wide (fig. S6B). Further studies should seek to determine whether depletion of Runx consensus sites in superenhancer candidates suppresses tumorigenesis in animal cancer models.

p53 deficiency and Myc excess, two major forces governing genesis and progression of most cancers in humans and mouse, are linked through Runx3 and mR1, providing a rationale for the targeting of Runx3 in cancer therapy. p53 is not amenable to pharmacological manipulation and has been widely deemed to be ‘undruggable’ (*40*). Instead of direct retrieval of p53, indirectly targeting Runx3 or mR1 using drugs that inhibit the interaction of Runx and Cbfβ (*23, 24*) as shown in this study (Fig. 4F), or PI polyamides targeting the consensus RUNX-binding sequences (*35*) could effectively achieve the same goal. The oncogenic Runx3–Myc axis is most likely to be critical for *p53*-deficient malignancies other than OS. Because this oncogenic axis is dormant in *p53*-proficient normal cells (Fig. 5H), it is an attractive and widely applicable target for anti-tumor pharmacotherapy in a variety of human cancers, in particular from the standpoint of avoiding side effects.

## Materials and Methods

### Mouse lines

Floxed mouse lines of *Runx1* (*41*), *Runx2* (*42*), *Runx3* (*43*), *p53* (*44*), and *Myc* (*45*) were described previously. The *Sp7/Osx*-Cre line (no.006361) was purchased from Jackson Laboratory. All mouse studies were performed in the C57BL/6 background.

A *Runx3(P1)*-deleted mouse line was generated that systemically lacked the 3’ region of the *Runx3(P1)* exon 1, which encodes the N-terminal region of Runx3(P1) (MASNSIFDSFPNYTPTFIR). The targeting vector harboring exon 1 and a *Loxp/FRT*-flanked Neomycin resistance gene (*Neo*) was electroporated into Bruce-4 ES cells (C57BL/6). Digested genomic DNA of targeted ES cells was subjected to Southern blotting using 5’ or 3’ alkaline phosphatase-labeled probes (386 bp or 536 bp), which were generated by PCR using primers 5’-TGTAATGCTCTGTCCTTTATCTGTG-3’ and 5’-CTGACTGTCTTCTGGTTAGAACGAT-3’, or 5’-GATGTAGCACCCAGTATCTCAAAGC-3’ and 5’-CAGATGCTAGACTTCCTTCTCAGTTA-3’, respectively, and mouse genome DNA as the template. To remove the *Loxp*-flanked *Neo*, the offspring (F1) was crossed with *CAG-*Cre transgenic mice. The *Runx3(P1)*^*Δ*^ and wild-type alleles of *Runx3(P1)*^*Δ/*+^ mice were detected by PCR using primers 5’-CTCAAACAGACAAAGGTCAGATGTA-3’ and 5’-ATGTTGAGCAGATTCTCATAGAGG-3’, yielding products of 205 bp and 642 bp, respectively.

An mR1-mutated mouse line was generated by homologous recombination. The targeting vector harboring a *BglII*-replaced mR1 and an *FRT*-flanked neomycin resistance gene (*Neo*) were electroporated into Bruce-4 ES cells. Digested genomic DNA of targeted ES cells was subjected to Southern blotting using 5’ or 3’ alkaline phosphatase-labeled probes (483 bp or 380 bp), which were generated by PCR using primers 5’-TTTGACATAGCTAGTGAGACAGCAG-3’ and 5’-CAGGAGGAGATATTTGAGGTTGTTA-3’, or 5’-CAGGATGCTTTCTGTGGATAGTAAT-3’ and 5’-CTCTGAACCTTTGTATGTCCTTGTT-3’, respectively, and mouse genome DNA as the template. To remove the *FRT*-flanked *Neo*, F1 was crossed with *CAG-*FLP transgenic mice. The mutant and wild-type alleles of *mR1*^m/+^ mice were detected by PCR using primers 5’-GGTTTAGAGTGTAGAAGGGAGGTGT-3’ and 5’-ATTGCTGACTTGGAGGAGAGAG-3’, yielding products of 188 bp and 110 bp, respectively.

Genome-edited mR1-, mR2-, or mR3-mutant mouse line was generated by intra-cytoplasmic injection of fertilized zygotes with Alt-R S.p. HiFi Cas9 Nuclease (Integrated DNA Technologies), an annealed pair of Alt-R CRISPR-Cas9 tracrRNA and Alt-R CRISPR-Cas9 crRNA (Integrated DNA Technologies), and ssDNA flanking the target site. DNA target sequences upstream of each PAM site (20 bases) and the corresponding ssDNA sequences are shown below.

mR1

crRNA: 5’-CTGCGTATATCAGTCACCGC-3’

ssDNA (150-mer):

5’-GGCCGCCCGGGACGTGCGTGACGCGGTCCAGGGTACATGGCGTATTGTGT GGAGCGAGGCAGCTGTTCCACCagatctGACTGATATACGCAGGGCAAGAACACA GTTCAGCCGAGCGCTGCGCCCGAACAACCGTACAGAAAGGGAAAG-3’

mR2

crRNA: 5’-AGGGTGATCAACCGCAGATG-3’

ssDNA (150-mer):

5’-TATACGTGGCAGTGAGTTGCTGAGCAATTTTAATAAAATTCCAGACATCGT TTTTCCTGCATAGACCTCATCagatctTGATCACCCTCTATCACTCCACACACTGAG CGGGGGCTCCTAGATAACTCATTCGTTCGTCCTTCCCCCTTT-3’

mR3

crRNA: 5’-TCGTTGGCTTCGCAACGCTG-3’

ssDNA (150-mer):

5’-CATCTTCCCAGAACCTGGAAACCCTGCAGCCCTGCCCCCATCCGACCTCC GCCCTCGTTGGCTTCGCAACGCagatctCTCTGTGGCCAGTAGAGGGCACACTTAC TTTACTTTCACAAATCCGAGAGCCACAACCCGGGTGGTGGGGGG-3’

For mouse genotyping, PCR products amplified using the following primers were digested with *Bgl*II, which replaced each of the native Runx binding sites.

mR1/mR2

5’-GTCGTTCTGGAAAGAATGTGC-3’

5’-GCCCAGTACTCCGGCTCC-3’

mR3

5’-GCAGCCTAAAAGAGTCATTTAAAGG-3’

5’-CCCAAGGCACTTAAAGAAACC-3’

All animal studies were reviewed and approved by the Animal Care and Use Committee of Nagasaki University Graduate School of Biomedical Sciences (no. 1603151292-14). Four mice were housed in each cage. Mice were reared in a pathogen-free environment on a 12 hour light cycle at 22 ± 2°C.

### BM-MSCs and mOS and human OS cells

All cells used in this study were maintained in F12/DMEM medium supplemented with 10% fetal bovine serum. For generation of BM-MSCs, BM cells were flushed from the femur of 3-month-old mice with F12/DMEM medium. Cd11b^-^ Cd45^-^ adherent BM cells, which were negatively selected using a magnetic cell sorting system (MACS) consisting of CD11b and CD45 MicroBeads and MS Columns (Miltenyi Biotec), were used as BM-MSCs. Similarly, for generation of mOS cells, adherent cells obtained from mouse OS tissues that were minced and collagenase I-digested were negatively selected using a MACS. Cd11b^-^ Cd45^-^ OS cells were cloned and used as mOS cells. hMSC (human mesenchymal stem cell)/NHOst (human osteoblast cell), U2OS/G292/Saos-2/SJSA1/MNNG-HOS/143B, MG-63/HOS/HuO9, and NOS1 cells were purchased from Lonza, ATCC, JCRB, and RIKEN, respectively.

### Immunoblotting and immunoprecipitation

Lysates of OS cells and MSCs for immunoblotting were prepared using a lysis buffer containing 9 M Urea, 2% Triton X-100, 2-mercaptoethanol, and proteinase/phosphatase inhibitors. OS tissues were homogenized using a FastPrep24 homogenizer (MP Bio). Immunoblotting was performed using the following antibodies: anti-Runx1 (in-house rabbit polyclonal antibody against Runx1 polypeptide MSEALPLGAPDGGAALAS), anti-Runx2 (D1H7; Cell Signaling Technology), anti-Runx3 (D6E2; Cell Signaling Technology), anti-Cbfβ (β-4E8; in-house mouse monoclonal antibody) (*46*), anti-VWRPY (Rp-6C2; in-house mouse monoclonal antibody), anti-Runx3(P1) (in-house rabbit polyclonal antibody against Runx3(P1) polypeptide SNSIFDSFPNYTPTFIRDP), anti-c-Myc (Y69, Abcam), anti-p53 (DO-1, MBL, 1C12, Cell Signaling Technology), anti-p21 (EPR3993, Abcam), anti-FLAG (M2; Sigma), and anti-β-actin (AC-15; Sigma).

Whole-cell extracts of mOS1-1 cells transiently expressing human wild-type p53 (pcDNA3.1-human p53WT), p53(R156P) (pcDNA3.1-p53R156P), or p53(R273H) (pcDNA3.1-p53R273H) and HEK293T cells transiently expressing human wild-type p53 and RUNX1, 2, or 3 (P1) (pcDNA3.1-RUNX1, 2, or 3-P1) were immunoprecipitated with anti-Runx3 (D6E2) and anti-p53 (DO-1) antibodies as appropriate. The immunoprecipitates from mOS1-1 or HEK293T cells were subjected to immunoblotting using anti-p53 (DO-1)/anti-Runx3 (R3-8C9) (*47*) antibodies or anti-VWRPY/anti-p53 (FL393; Santa Cruz) antibodies, respectively. The anti-VWRPY antibody was evaluated using whole-cell extracts of HEK293T cells transiently expressing FLAG-tagged full-length Runxs(P1) (pFLAG-CMV-4-Runx1, 2 or 3-P1) or FLAG-tagged full-length Runxs(P1) without VWRPY (pFLAG-CMV-4-Runx1, 2 or 3-P1 ΔVWRPY) and an anti-FLAG antibody. Immunoprecipitates of nuclear extracts (NE) of ST2 cells generated using anti-Runx1, anti-Runx2 (D1H7), or anti-Runx3 (D6E2) antibodies or normal rabbit IgG were subjected to immunoblotting with the anti-VWRPY antibody to confirm the positions of endogenous Runx1, 2, and 3 proteins on the western blotting. To detect endogenous protein interactions, immunoprecipitates of NE of ST2 cells generated using anti-Runx2 (D1H7) or anti-Runx3 (D6E2) antibodies or normal rabbit IgG were subjected to immunoblotting using anti-Runx2 (R2-8G5; MBL), anti-Runx3(R3-8C9), or anti-p53 (1C12) antibodies.

### shRNA-knockdown and transplantation

Knockdown of endogenous gene expression was performed using the RNAi-Ready pSIREN-RetroQ Vector (Clontech). Cells were retrovirally transfected and selected with puromycin. Resistant cells were used for subsequent assays without cloning. shRNA sequences targeting *Runx1, Runx2*/*RUNX2, Runx3*/*RUNX3*, or *Myc*/*MYC* are shown below:

Runx1-1: 5’-AAGACATCGGCAGAAACTAGATGAT-3’

Runx1-2: 5’-AGCTTCACTCTGACCATCA-3’

Runx2/RUNX2-1: 5’-GTTGCAACTGTAAATTGAA-3’

Runx2/RUNX2-2: 5’-GGACTGTGGTTACCGTCAT-3’

Runx3-1: 5’-GGAGCCATATCTCTCTTTC-3’

Runx3-2: 5’-GCATCTCAGTCAAGCATCT-3’

RUNX3-1: 5’-GCCCCAGAGAAGATGAGTCTAT-3’

RUNX3-2: 5’-TCAGTAGTGGGTACCAATCTT-3’

Myc-1: 5’-GAACATCATCATCCAGGAC-3’

MYC-1: 5’-AACAGAAATGTCCTGAGCAAT-3’

Myc/MYC-2: 5’-ACATCATCATCCAGGACTG-3’

Transplantation (xenografts and allografts) was performed by subcutaneous injection of human OS (5 × 10^6^) or mOS (2.5 × 10^6^) cells into BALB/c-*nu/nu* mice (nude mice). Tumorigenicity of OS cells was assessed based on tumor weight 4 weeks after inoculation.

## qRT-PCR

Total RNA was extracted using the NucleoSpin RNA kit (Macherey-Nagel) and reverse-transcribed into cDNA using the ReverTra Ace qPCR RT Master Mix (Toyobo). Real-time quantitative PCR reactions were performed on a 7300 Real-time PCR system (ABI) using THUNDERBIRD Probe or SYBR qPCR Mix (Toyobo). The following TaqMan probes (Applied Biosystems) or primer sets were used:

*Runx1*: Mm00486762_m1

*Runx2*: Mm03003491_m1

*Runx3*: Mm00490666_m1 or 5’-ATGAAGAACCAAGTGGCCAGG-3’ and 5’-CTTGATGGCTCGGTGGTAGG-3’

*Myc*: Mm00487803_m1 or 5’-GTGCTGCATGAGGAGACACC and 5’-CACAGACACCACATCAATTTCTTCC-3’

*β-actin*: Mm00607939_s1 or 5’-CATCCGTAAAGACCTCTATGCCAAC-3’ and 5’-ATGGAGCCACCGATCCACA-3’

*Pvt1*: 5’-GTGAAGCGTTGACTTAAGAGATGC-3’ and 5’-TGCAGAACTCAGCTGTCTTATAGG-3’

### Digital PCR

Droplet generation and transfer of emulsified samples to PCR plates was performed using a QX200 Droplet Generator (Bio-Rad). Plates were incubated using a thermal cycler with the following conditions: 95°C for 5 min, 40 cycles of 95°C for 30 sec and 60°C for 1 min, 4°C for 5 min, and 90°C for 5 min. Plates were then read on a QX200 Droplet Reader (Bio-Rad), and the data were analyzed using the QuantaSoft Software (Bio-Rad). The following primer pairs were used:

*β-actin*: 5’-CATCCGTAAAGACCTCTATGCCAAC-3’ and 5’-ATGGAGCCACCGATCCACA-3’

*Runx3(P1)*: 5’-CAAAACAGCAGCCAACCAAGTGG-3’ and 5’-GGTTGGTGTATAGTTGGGGAAGG-3’

*Runx3(P2)*: 5’-TGACGGCCGCGGCATGCGTATTCCC-3’ and 5’-GCTGTTCTCGCCCATCTTGCC-3’

### RNA-seq

Two mouse calvariae (OB) were isolated from newborn mice. Seven mouse OS tissues were isolated from seven individual *OS* mice upon osteosarcoma development. Calvariae and OS tissues were homogenized using a FastPrep24 homogenizer (MP Bio). RNA was extracted from homogenates of these tissue samples using the NucleoSpin RNA kit. RNA quality was assessed on a Bioanalyzer (Agilent). Libraries were prepared using the SureSelect XT RNA Direct System (Agilent) after exome capture was performed using the probe set accompanying the SureSelect XT Mouse All Exon Kit (mm9). Next-generation sequencing (NGS) libraries were sequenced on an MGI-SEQ platform, with an average of 52 million reads and an average mapping rate of 97%. RNA counts were quantified using Kallisto(*48*) by pseudo-aligning FASTQ reads to the mouse genome (mm10).

RNA-seq data and the corresponding clinical information from 89 primary osteosarcoma patients (hOS) were obtained from the TARGET project (https://ocg.cancer.gov/programs/target) under accession number phs000468 on NCBI dbGaP. Of those, 83 datasets possessing both sequencing and clinical information were used for downstream analyses. RNA-seq datasets from two human osteoblast (hOB) samples were derived from the ENCODE project under accession number GSE78608. RNA counts were quantified using Kallisto(*48*) by pseudo-aligning FASTQ reads to the human genome (hg36).

In each of the four groups [mouse (m) OB, mOS, human (h) OB, and hOS], genes were excluded from subsequent analyses if their average expression counts were zero. Principal component analysis (PCA) was performed using the prcomp function of R. Differential expression analysis of osteosarcoma (mOS, hOS) vs. control osteoblasts (mOB, hOB) was performed using TMM/edgeR in the TCC platform(*49*), and the results were used to draw both MA plots and heatmaps. The 794 human genes associated with the Gene Ontology term “DNA-templated transcription” (GO:0000977; https://www.ebi.ac.uk/QuickGO/term/GO:0000977) and the 741 corresponding mouse genes were considered as transcription factors.

Prior to correlation and survival analyses, human gene counts were normalized against the housekeeping gene *ACTB*, which encodes beta-actin. Correlation analysis was performed using Spearman’s correlation. Survival analysis was performed, and the hazard ratio was calculated with the Cox proportional hazard model after patients were stratified into low- and high-expressing groups based on the median value for each gene.

### Microarrays

Gene expression profiles of BM-MSCs derived from 3-month-old *OS* (n=4) and *OS*; *Runx3(P1)*^Δ/Δ^ (n=5) mice were assessed by oligonucleotide microarray analysis using the SurePrint G3 Mouse GE Microarray kit 8×60k (Agilent Technologies). Total RNA samples were used for the microarray analyses.

### ChIP-qPCR and ChIP-seq

ChIP-seq experiments were performed using the SimpleChIP Enzymatic Chromatin IP kit with magnetic beads (Cell Signaling Technology). Briefly, 6 million cells were cross-linked with 1% formaldehyde for 10 min at room temperature. After permeabilization, cross-linked cells were digested with micrococcal nuclease and immunoprecipitated with isotype control, anti-Runx3 (D6E2; Cell Signaling Technology), anti-H3K27ac (D5E4; Cell Signaling Technology), or anti-RNAPII subunit B1 (Rpb1) NTD (D8L4Y; Cell Signaling Technology) antibodies. Immunoprecipitated products were isolated with Protein G Magnetic Beads (Cell Signaling Technology) and subjected to reverse cross-linking. The DNA was subjected to quantitative PCR using the following primer pairs.

ChIP-qPCR primers

Mouse gene desert: 5’-ACCAAGCACAGAAAAGGTTCAAAC-3’ and 5’-TCCAGATGCTGAGAGAAAAACAAC-3’

mR1: 5’-GCCTTAGAGAGACGCCTGGC-3’ and 5’-CCGCAGGTGGAACAGCTGC-3’ mR2: 5’-GTGGCAGTGAGTTGCTGAGC-3’ and

5’-GAAAGGGGGAAGGACGAACG-3’

mR3: 5’-CCCTCGTTGGCTTCGCAACG-3’ and 5’-ACCCGGGTTGTGGCTCTCG-3’

*Spp1* promoter: 5’-AATGACATCGTTCATCAGTAATGC-3’ and 5’-CATGAGGTTTTTGCCACTACC-3’

*Gapdh* promoter: 5’-GGACTGCCTGGTGTCCTTCG-3’ and 5’-TCACCCGTTCACACCGACC-3’

*Runx3* P1 promoter: 5’-ACACTGGGAAGGTCTGGTCC-3’ and 5’-AGGAACCAACCAGCTCCTCG-3’

*Myc* P1 promoter: 5’-CCCTTTATATTCCGGGGGTCTGC-3’ and 5’-CGAAGCCCTGCCCTTCAGG-3’

*Myc* P2 promoter: 5’-TCGCAGTATAAAAGAAGCTTTTCGG-3’ and 5’-CACACACGGCTCTTCCAACC-3’

*Myc* P3 promoter: 5’-CAGACAGCCACGACGATGC-3’ and 5’-CTTCCTCGTCGCAGATGAAATAGG-3’

*Myc* −0.1k: 5’-CCTCCCGAGTTCCCAAAGC-3’ and 5’-CGGGGATTAGCCAGAGAATCTC-3’

*Myc* +0.3k: 5’-GCTTTGCCTCCGAGCCTGC-3’ and 5’-GCAATGGGCAAAGTTTCCCAGC-3’

*Myc* +0.5k: 5’-GTCTATTTGGGGACAGTGTTCTCTG-3’ and 5’-GGGTTTCCAACGCCCAAAGG-3’

*Myc* Exon2: 5’-GTACCTCGTCCGATTCCACG-3’ and 5’-CAGCACTAGGGGCTCAGG-3’

*Myc* Exon3: 5’-TCTCGTGAGAGTAAGGAGAACG-3’ and 5’-CTGAAGCTTACAGTCCCAAAGC-3’

Human gene desert: 5’-TGAGCATTCCAGTGATTTATTG-3’ and 5’-AAGCAGGTAAAGGTCCATATTTC-3’

MR1: 5’-CCTGCGATGATTTATACTCACAGG-3’ and 5’-AAACCCTCTCCCTTTCTCTGC-3’

MR2: 5’-GCACGGAAGTAATACTCCTCTCC-3’ and 5’-GACGTTTAATTCCTTTCCAGGTCC-3’

MR3: 5’-GGGTGATGTTCATTAGCAGTGG-3’ and 5’-GAAGAAAGAGGAGTTACTGGAGG-3’

*SPP1* promoter: 5’-TTTAACTGTAGATTGTGTGTGTGC-3’ and

5’-ATTGTGTCATGAGGTTTTCTGC-3’

*RUNX3* P1 promoter: 5’-CACTGGGAAGGCCTGGTCC-3’ and

5’-GCACCAGGAGCCAACCAGC-3’

For high-throughput sequencing, libraries were prepared using the NEBNext Ultra II DNA Library Prep Kit for Illumina, and then purified with AMPure XP beads. Libraries were sequenced on the Illumina HiSeq platform.

### ATAC-seq

ATAC-seq experiments were performed as described(*50*). Briefly, 50,000 cells were washed with PBS and lysed in lysis buffer (10 mM Tris-HCl, 10 mM NaCl, 3 mM MgCl_2_, 0.1% Igepal CA-630). Transposed DNA fragments were generated using the Tagment DNA TDE1 Enzyme and Buffer Small Kit (Illumina), and amplified by PCR with an additional two cycles relative to the original protocol(*50*) using NEBNext High-Fidelity 2× PCR Master Mix (New England Biolabs). ATAC-seq libraries were sequenced on the Illumina HiSeq platform.

### Analysis of ChIP-seq and ATAC-seq

Reads were trimmed of adapter sequences using fastp (*51*) and aligned to the mouse genome (mm10) using bowtie2 (*52*). After removal of PCR duplicates with Picard MarkDuplicates (http://broadinstitute.github.io/picard), peaks were called using the findPeaks function of HOMER (*53*) (http://homer.ucsd.edu/homer) with the input set as a control. Correlations between ChIP-seq and ATAC-seq samples were calculated using the plotCorrelation function of deepTools (*54*). De novo motif prediction was performed using the findMotifs functions of HOMER with default settings. All peaks of ATAC-seq and ChIP-seq samples were merged and partitioned into three clusters according to the scores calculated by the mergePeaks feature of HOMER (*53*). Along the positions of the merged peaks, heatmaps were drawn using the plotHeatmap function of deepTools. The 1624 genes nearest the peaks detected in the strongly co-occupied cluster were considered significant. Topologically associating domain (TAD) boundaries for *Myc* were determined by referring to previously used (*55*).

### Histological analysis

Tissues were fixed with 4% paraformaldehyde in PBS, decalcified in Osteosoft (Merck Millipore) at room temperature for 10 days, embedded in paraffin, and cut into 4 µm sections. Anti-Myc (Y69, Abcam), anti-Runx2 (Cell Signaling Technology; D1H7), and anti-Runx3 rabbit (Cell Signaling Technology; D6E2) or mouse (R3-8C9) antibodies were used for immunodetection on rehydrated sections pretreated with Target Retrieval Solution (DAKO). The Envision™+ system (HRP/DAB) (DAKO) or Alexa Fluor (488 or 594; Invitrogen) were used for visualization.

### Epigenome editing

HEK293T cells were cotransfected with individual sgRNA-dCas9-KRAB lentiviral expression plasmids (#71236; Addgene), the second-generation packaging plasmid psPAX2 (#12260; Addgene), and the envelope plasmid pMD2.G (#12259; Addgene) by a standard lipofection method. After filtration with a 0.45-µm filter, c, conditioned medium containing lentivirus was used for transduction. Transduced cells were selected with puromycin. Expression of FLAG-tagged dCas-KRAB in mOS1-1 cells was immunodetected using an anti-FLAG antibody (M2). sgRNA sequences were as follows:

Scrambled: 5’-TGGTTTACATGTCGACTAAC-3’

Myc-740k: 5’-TCCATATAAGTGACTGAATG-3’

Myc-450k: 5’-GCACCTAGGCATCCCACTTG-3’

Myc mR3: 5’-TCGTTGGCTTCGCAACGCTG-3’

Myc mR2: 5’-ACCGCAGATGAGGTCTATGC-3’

Myc mR1: 5’-CTGCGTATATCAGTCACCGC-3’

Myc TSS: 5’-CGCTCCGGGGCGACCTAAGA-3’

Myc +300k: 5’-CCTTAAACTCAACCCTCCAG-3’

Myc +340k: 5’-CAGAACCAATAAGAGCGGTG-3’

Myc +710k: 5’-ATAAGTTCAGAACATTCTGT-3’

Myc +1.1M: 5’-GTGCCAGCAGATTCACCTCT-3’

Myc N-Me: 5’-GCTGATGATTTCAGTCAATT-3’

Myc BDME-E3: 5’-ACACAGTGTGGTTCCTTTTC-3’

### Genome editing of OS cells

Mouse mR1 and mR3 were mutated by CRISPR-based genome editing, using the plasmid PX459 (#62988; Addgene). sgRNA sequences were as follows: Scrambled (5’-TGGTTTACATGTCGACTAAC-3’), mR1 (5’-CTGCGTATATCAGTCACCGC-3’), or mR3 (5’-TCGTTGGCTTCGCAACGCTG-3’). Cells were electroporated with the individual plasmids using the Neon Transfection System (Invitrogen).

Human MR1 was mutated by TALEN-based genome editing, using the Platinum Gate TALEN Kit (#1000000043; Addgene). The sequence 5’-TTATACTCACAGGACAAggatgcggtttgtcaaaCAGTACTGCTACGGAGG was used as the target; capital letters indicate sequences recognized by the two corresponding TALEN vectors constructed using the kit. Cells were electroporated with the pair using the Neon Transfection System. Clones harboring the intended mutation were identified by sequencing.

### Stable and inducible expression of exogenous genes

Cells were retrovirally transfected with pMSCV-vector (Clontech/Takara), pMSCV-Runx3(P1), pMSCV-RUNX3(P1), or pMSCV-human-p53WT and selected using puromycin. Resistant cells were cloned. Expression of human p53WT, p53(R156P), or p53(R273H) was induced in mOS cells using Retro-X Tet-One Inducible Expression System (Clontech/Takara). mOS cells were retrovirally transfected and selected with puromycin. Resistant cells were cloned and treated with or without 100 ng/ml doxycycline (Dox) for the indicated times.

### Luciferase reporter assays

Runx(wt)- or Runx(mt)-Luc vectors harbored 6×Runx wild-type (wt) or mutant (mt) binding sites (wt: 5’-TGCGGTTGCGGTTGCGGTgtcTGCGGTTGCGGTTGCGGT-3’; mt: 5’-TGCCCTTGCCCTTGCCCTgtcTGCCCTTGCCCTTGCCCT-3’) between the *Xho*I and *Hin*dIII sites of the pGL4.10[luc2] vector (Promega), respectively. G292 cells were co-transfected with Runx(wt)- or Runx(mt)-Luc and an internal control, pRL-SV40 (Promega), in combination with pcDNA3.1-mock, -human p53WT, -p53(R156P), -p53(R273H), -RUNX2/Runx2(P1), or -RUNX3/Runx3(P1). Transfections were performed using the X-tremeGENE HP DNA Transfection Reagent (Roche). Forty-eight hours after transfection, luciferase activity was measured using the dual-luciferase reporter assay system (Promega).

### EMSA

EMSA was performed using the LightShift Chemiluminescent EMSA Kit and Chemiluminescent Nucleic Acid Detection Module (Thermo Scientific). Each binding reaction (10 μl) contained 50 ng/μl poly (dI-dC), 75 fmol labeled probe, and NE in the buffer supplied in the kit. NE was prepared from ST2, MC3T3-E1, and mOS cells, as well as from 293T cells expressing p53 (pcDNA3.1-human p53) or harboring pcDNA3.1, using the NE-PER Nuclear and Cytoplasmic Extraction Reagents (Thermo Fisher Scientific). NE and a labeled DNA probe with or without unlabeled probes were incubated at room temperature for 20 min and resolved on 5% polyacrylamide gels in 0.5x TBE buffer. For super-shift analysis, anti-Runx1, anti-Runx2 (D1H7), or anti-Runx3 (D6E2) antibodies or rabbit normal IgG were added after the binding reaction and incubated at room temperature for an additional 10 minutes. NE of p53- or mock-293T cells was incubated on ice for 10 min with NE of mOS1-1 cells, and then for 20 min at room temperature with labeled DNA probe. The following oligonucleotide probes were used: 5’-biotinylated labeled or unlabeled wild-type (wt) mR1, 5’-TTCCACCTGCGGTGACTGAT-3’; 5’-biotinylated labeled T7 or T13, 5’-TTCCACGCGGTGACTGAT-3’ or 5’-TTCCACCGGTGACTGAT-3’, respectively; and unlabeled mutated (mt) mR1, 5’-TTCCACCTGCCGTGACTGAT-3’.

### Administration of Runx inhibitors

*OS* mice that developed OS in the lower limbs, the most frequent site, were selected, and administration of the inhibitors via intraperitoneal injection was initiated when the onset of OS was visually confirmed (i.e., when the tumor was around 3 mm in diameter). Ro5-3335 or AI-10-104 were administered at 5 mg/kg in 100 μl of 50% DMSO in PBS or 1 mg/kg in 100 μl of 10% DMSO in PBS, respectively, once every 3 or 4 days (twice a week) for 10 weeks after OS onset.

### Statistics

All quantitative data are expressed as means ± SD. Differences between groups were calculated by unpaired two-tailed Student’s t-test for two groups or by one-way analysis of variance (ANOVA) for more than two groups. All analysis were performed in Prism 8 (GraphPad software). Survival was analyzed by the Kaplan–Meier method and compared by the log-rank test using the same software. *p*<0.05 denotes significance.

## Acknowledgments

We thank G. Huang for critical advice on the study; A. Berns and F. W. Alt for providing the *p53* and *Myc* flox mouse lines, respectively; T. Kishino for generating genome-edited mouse lines; and all members of the Biomedical Research Center, Nagasaki University for maintaining mouse lines.

## Funding

This work was supported by KAKENHI/Japan Society for the Promotion of Science (JSPS) grants 26290040 (K.I.), 18H02972 (K.I.), and 19K22724 (K.I.); by the Funding Program for Next Generation World-Leading Researchers LS097 (K.I.); and by the JSPS Research Fellowship for Young Scientists (Y.D.).

## Author contributions

K.I. initiated the study. Y.D. and K.I. designed the experiments. S.O., Y.D., T.U., T.I., S.K., K.O., J.T., and K.I conducted the experiments. Y.D. performed bioinformatic analyses. I.T. and T.K. generated and provided animal materials. S.O, Y.D., and K.I. wrote the manuscript. M.U. and T.K. coordinated the project. K.I supervised the study.

## Competing interests

The authors declare no competing interests.

## Data and materials availability

All data are available in the main text or the supplementary materials. The accession numbers for the ChIP-seq/ATAC-seq and RNA-seq data generated in this study are DRA009517 and DRA011168, respectively.

## Supplementary Materials

Figures S1–S13

Tables S1–S2

## Notes

### Competing Interest Statement

The authors have declared no competing interest.

