## Supplementary Materials for "The oncogenic Runx3–Myc axis defines *p53*-deficient osteosarcomagenesis"

**This file includes:**

Figs. S1 to S13
Tables S1 and S2

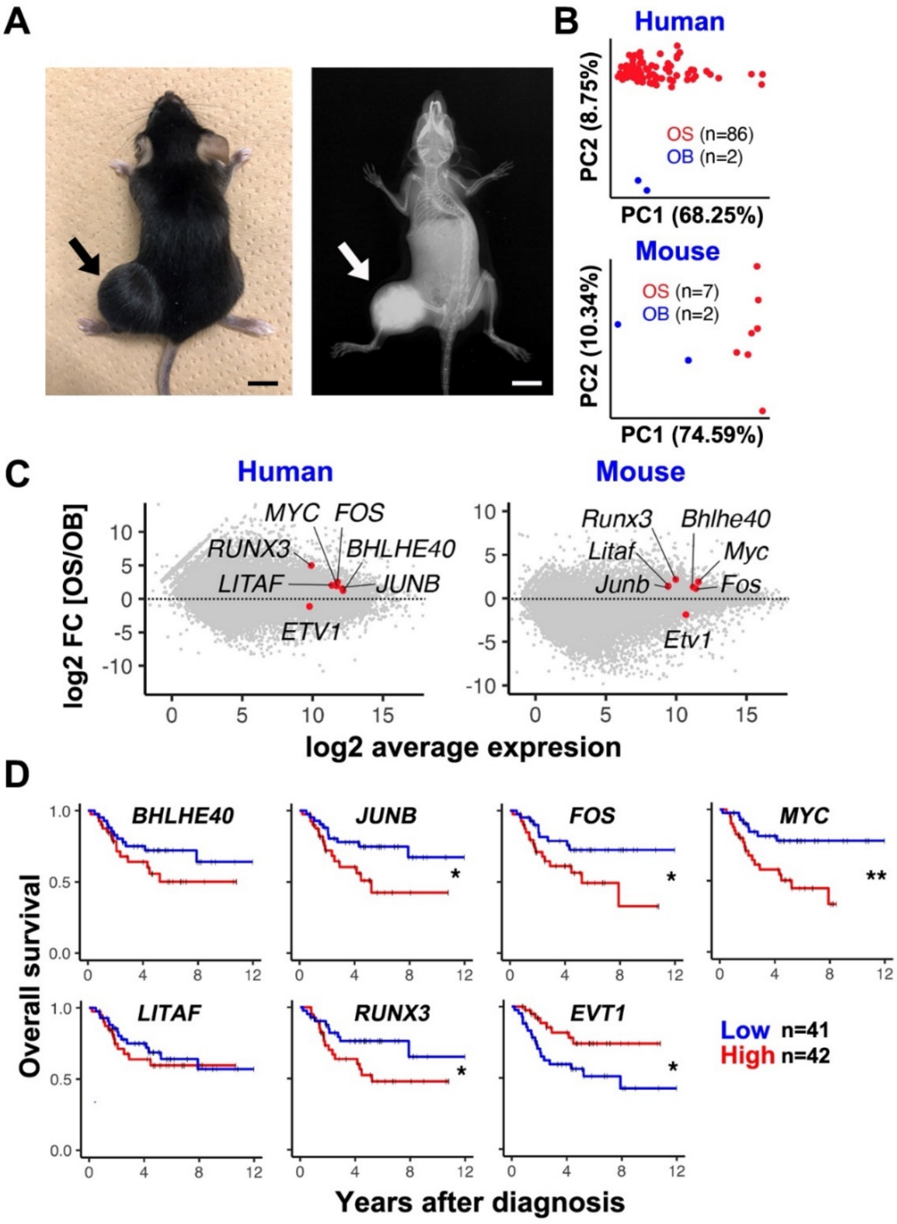


S Figure 1

**Fig. S1 Up- and down-regulated transcription factors in *p53*-deficient osteosarcomagenesis in human and mouse.**

(**A**) OS development in the tibia of a representative osteoprogenitor-specific *p53* knockout mouse (*Osterix/Sp7*-Cre; $p53^{fl/fl}$). Arrows indicate the tumor. *Osterix*-Cre is expressed in BM-MSCs of adult mice as well (*56*). Right panel: radiograph. A scale bar = 1 cm. (**B**) Principal component analysis (PCA) of RNA-seq data comparing OS tissues (OS) and normal osteoblasts (OB) in human and mouse. (**C**) MA plot presentation of RNA-seq data comparing OS and OB. The seven TFs (Fig. 1, A and B) are indicated by red dots. (**D**) Prognostic efficacy of the seven transcription factors (Fig. 1, A and B) as determined by Kaplan–Meier analysis of survival in the OS patients in the TARGET cohort. $^{**}p < 0.01$; $^{*}p < 0.05$.

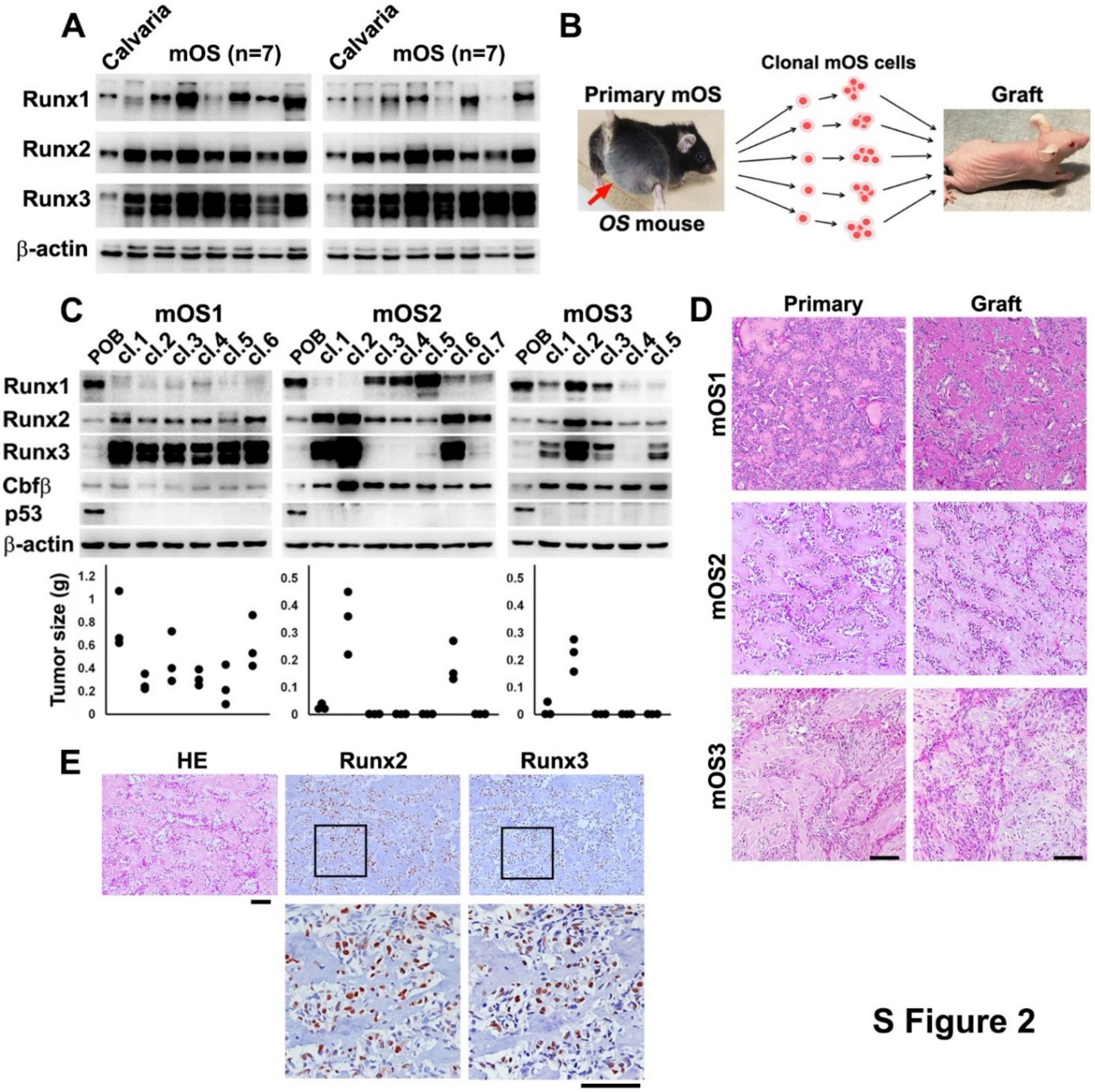


**Fig. S2 Runx3 is required for tumorigenicity of *p53*-deficient OS.**

(**A**) Levels of the indicated proteins in OS (mOS) in 14 individual *OS* mice and newborn calvaria, as determined by western blotting. (**B**) Tumorigenicity of clonal mOS cells isolated from mOS (arrow) in individual *OS* mice (left) was evaluated by allograft in nude mice (BALB/c *nu/nu* mice; right). (**C**) Levels of the indicated proteins in clonal OS cells isolated from OS (mOS1~3) in three individual *OS* mice, and primary osteoblasts (POB) derived from newborn calvaria, as determined by western blotting (upper). The tumorigenicity of each clone was evaluated by allograft (lower; n=3). Clone 1 of mOS1 and clone 2 of mOS2 were used as mOS1-1 and mOS2-2 cells, respectively, in this study. (**D**) Strong histological similarity of primary OS (mOS1–3) in three individual *OS* mice (Primary; left panels) and allograft OS of mOS1 clone 1, mOS2 clone 2, and mOS3 clone 2 (grafts; top right, middle right, and bottom right, respectively). (**E**) Representative histology of OS in *OS* mice (HE; left) and immunodetection of Runx2 and Runx3 (center and right). Serial sections were used. Boxed regions are enlarged below. Counter staining was done with hematoxylin. Scale bars = 100 μm.

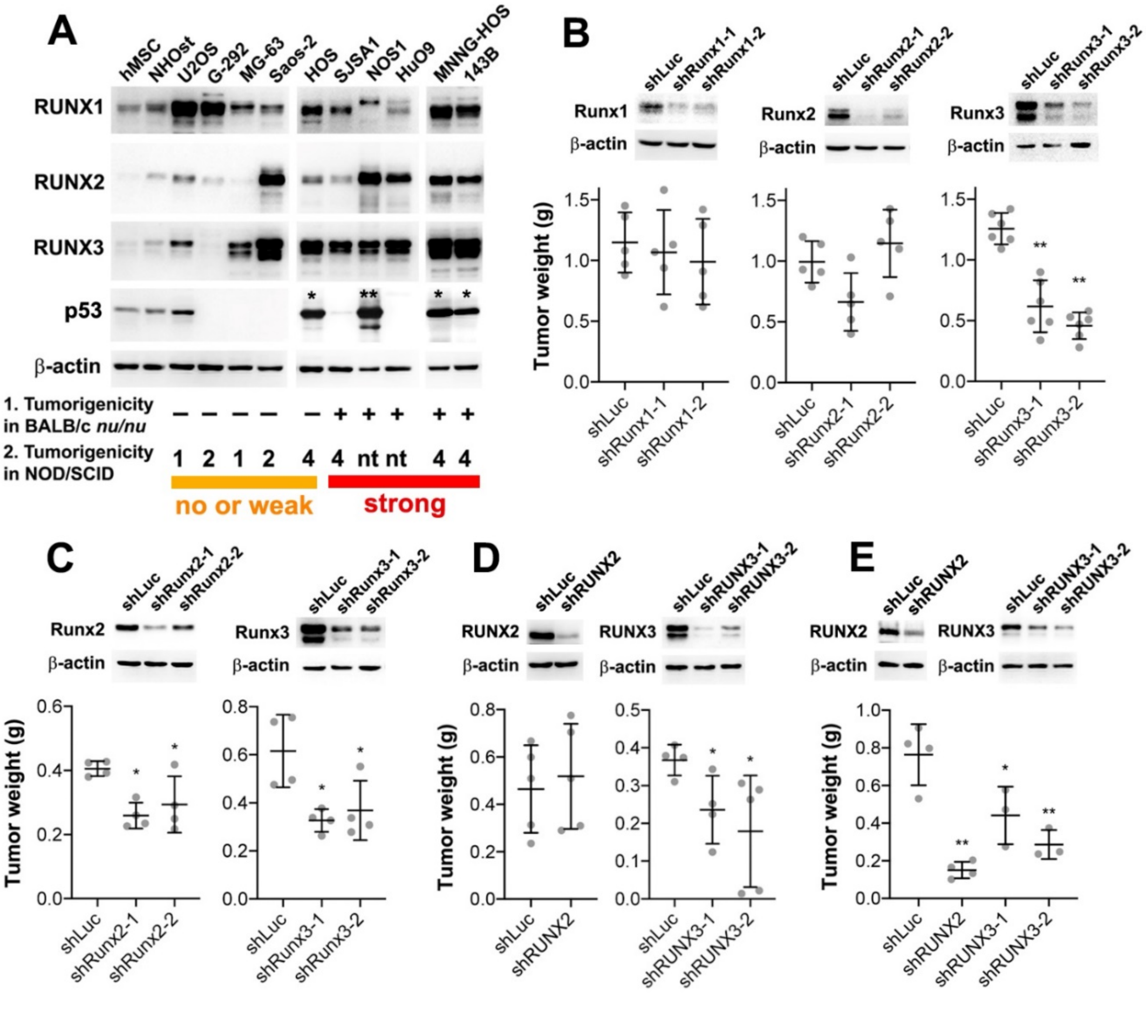


S Figure 3

**Fig. S3 RUNX3/Runx3 is required for tumorigenicity of human and mouse *p53*-deficient OS.**

(**A**) Expression profile of the indicated proteins in human OS cell lines, normal human MSC (hMSC), and osteoblast (NHOst), as determined by western blotting. Tumorigenicity of the human OS cell line was classified based on previously published results [1 (*57*) and 2 (*58*)]. NOS1 and HuO9 were also tumorigenic in xenografts using BALB/c *nu/nu* mice as reported (*59*) and shown in (**E**), respectively. Mutated p53s are indicated by asterisks: *R156P and **R273H (fig. S11E). nt, not tested. (**B**) Effects of knockdown of Runx1, Runx2, or Runx3 on tumorigenicity of mOS1-1 cells (mOS1 cl.1 cells shown in fig. S2C). Levels of the indicated proteins in cells, as determined by western blotting (upper). Tumorigenicity was evaluated by allograft (lower). (**C**) Effects of knockdown of Runx2 or Runx3 on tumorigenicity of mOS2-2 cells (mOS2 cl.2 cells shown in fig. S2C). Levels of the indicated proteins in the cells were determined by western blotting (upper). Tumorigenicity was evaluated by allograft (lower). (**D** and **E**) Effects of knockdown of RUNX2 or RUNX3 on tumorigenicity of SJSA1 (**D**) and HuO9 (**E**) cells. Levels of the indicated proteins in the cells were determined by western blotting (upper). Tumorigenicity was evaluated by xenograft (lower). $^{**}p < 0.01$; $^{*}p < 0.05$.

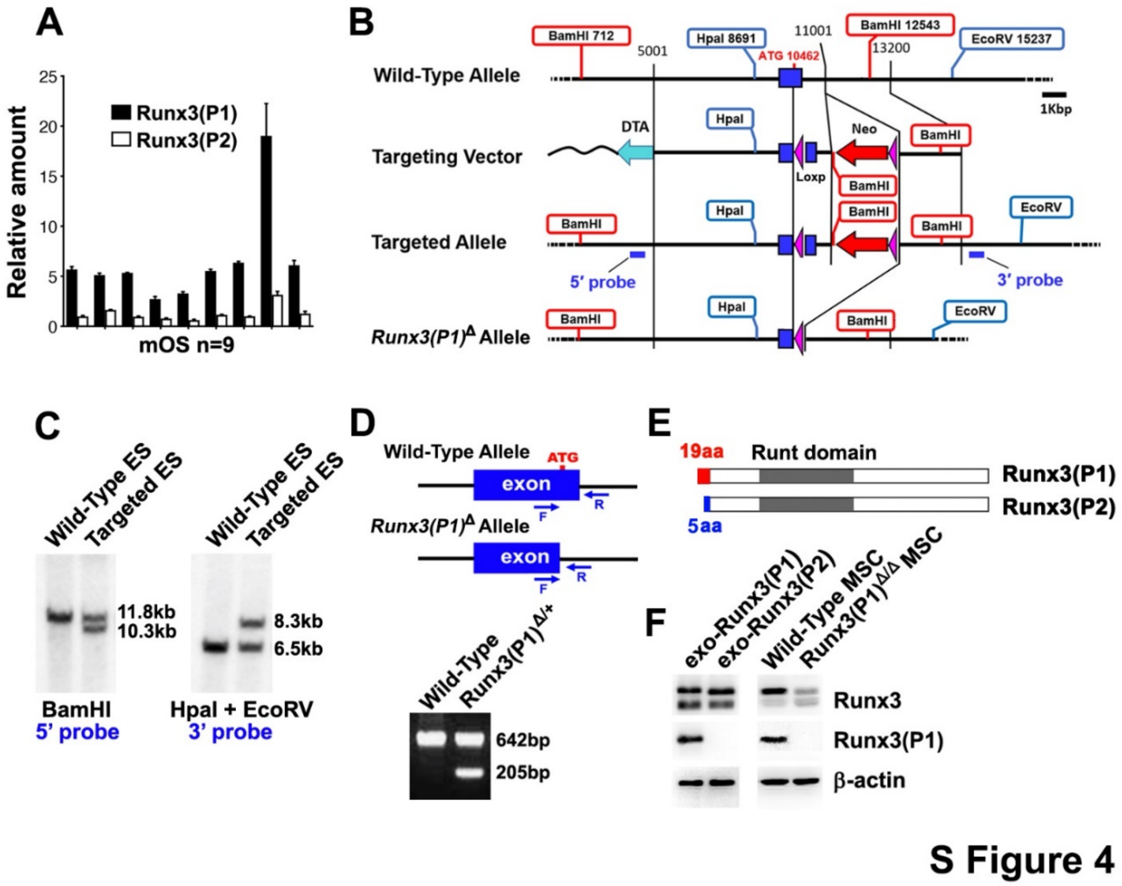


**Fig. S4 Generation of *Runx3(P1)*-deleted mice.**

(**A**) Levels of mRNA of *Runx3(P1)* and *Runx3(P2)* in nine OS (mOS) developed in nine individuals of *OS* mice, as determined by digital PCR (n=3). (**B**) Targeting vector for homologous recombination and strategy for generating the *Runx3(P1)*-deleted mouse line. (**C**) Southern blot analyses of genomic DNA of wild-type and targeted ES cells digested with *BamHI* (left) or *HpaI* and *EcoRV* (right) using 5' or 3' probes shown in (**B**), respectively. (**D**) Genotyping PCR for detection of wild-type and *Runx3(P1)*$^{\Delta}$ alleles using F and R primers. (**E**) Structure of Runx3(P1) and Runx3(P2) proteins. The distinct N-terminal amino acids of Runx3 (P1) and Runx3 (P2) are (M)ASNSIFDSFPNYTPTFIR (19 aa) and (M)RIPV (5 aa), respectively. (**F**) Runx3(P1) was not expressed in BM-MSCs of *Runx3(P1)*$^{\Delta/\Delta}$ mice, as determined by western blotting with an anti-Runx3(P1)-specific antibody (right). Whole-cell extracts of HEK293T cells exogenously expressing Runx3(P1) or Runx3(P2) were subjected to western blotting (left).

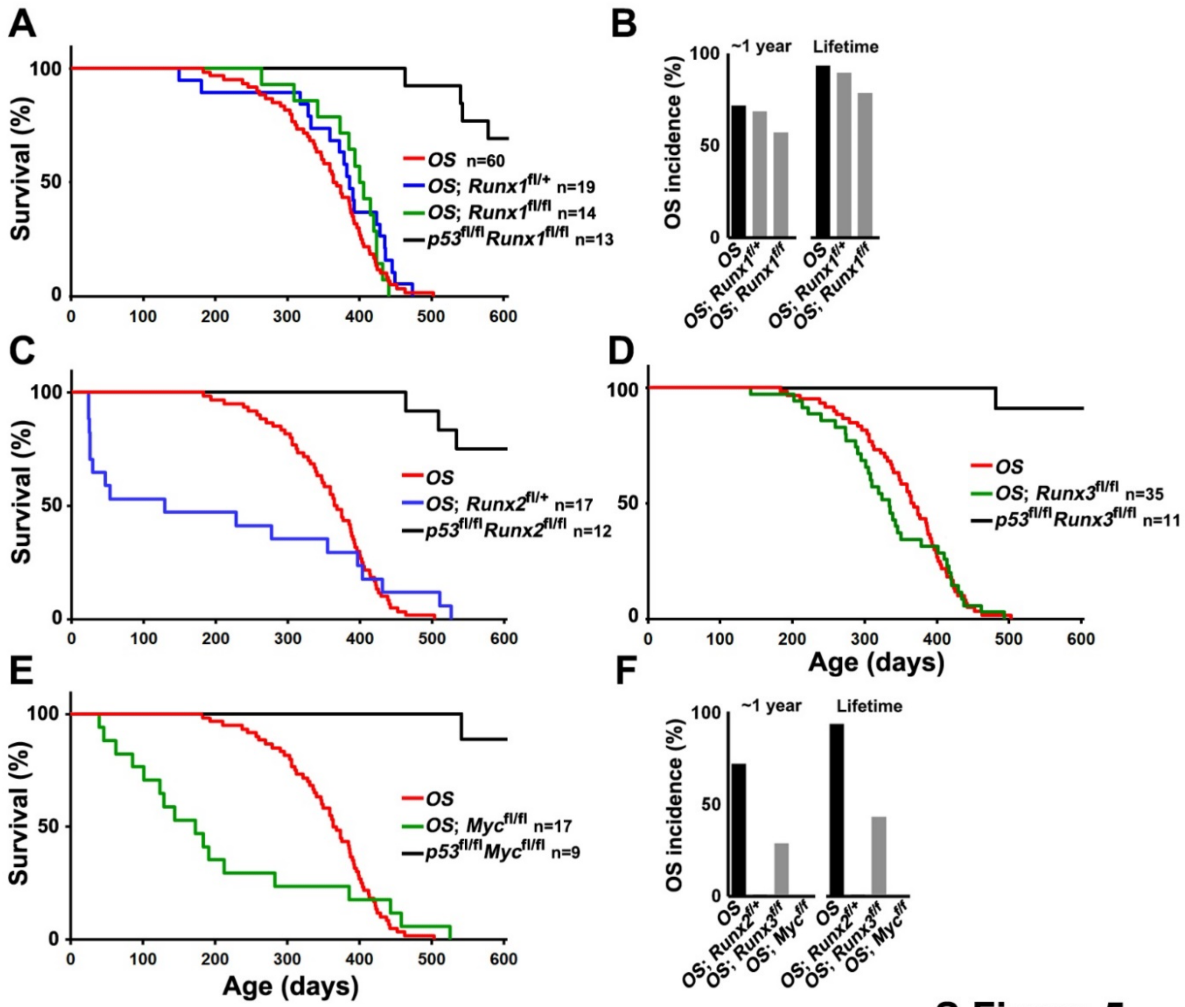


S Figure 5

**Fig. S5 Effects of heterozygous knockout of *Runx1* or *Runx2* and homozygous knockout of *Runx1*, *Runx3*, or *Myc* on *OS* mice.**

(**A**) Survival of the indicated mouse lines. *p53*$^{\text{fl/fl}}$*Runx1*$^{\text{fl/fl}}$ line is a Cre-free control. (**B**) OS incidence in the mouse lines shown in (**A**) within 1 year and throughout the lifespan. (**C**) Survival of the indicated mouse lines. *p53*$^{\text{fl/fl}}$*Runx2*$^{\text{fl/fl}}$ line is a Cre-free control. (**D**) Survival of the indicated mouse lines. *p53*$^{\text{fl/fl}}$*Runx3*$^{\text{fl/fl}}$ line is a Cre-free control. (**E**) Survival of the indicated mouse lines. *p53*$^{\text{fl/fl}}$*Myc*$^{\text{fl/fl}}$ line is a Cre-free control. (**F**) OS incidence in the mouse lines shown in (**C**) to (**E**) within 1 year and throughout the lifespan. In *OS*; *Runx2*$^{\text{fl/+}}$ and *OS*; *Myc*$^{\text{fl/fl}}$ mice, no OS was observed. The results of *OS* mice are identical to those in Fig. 1, D and E (**A** to **F**).

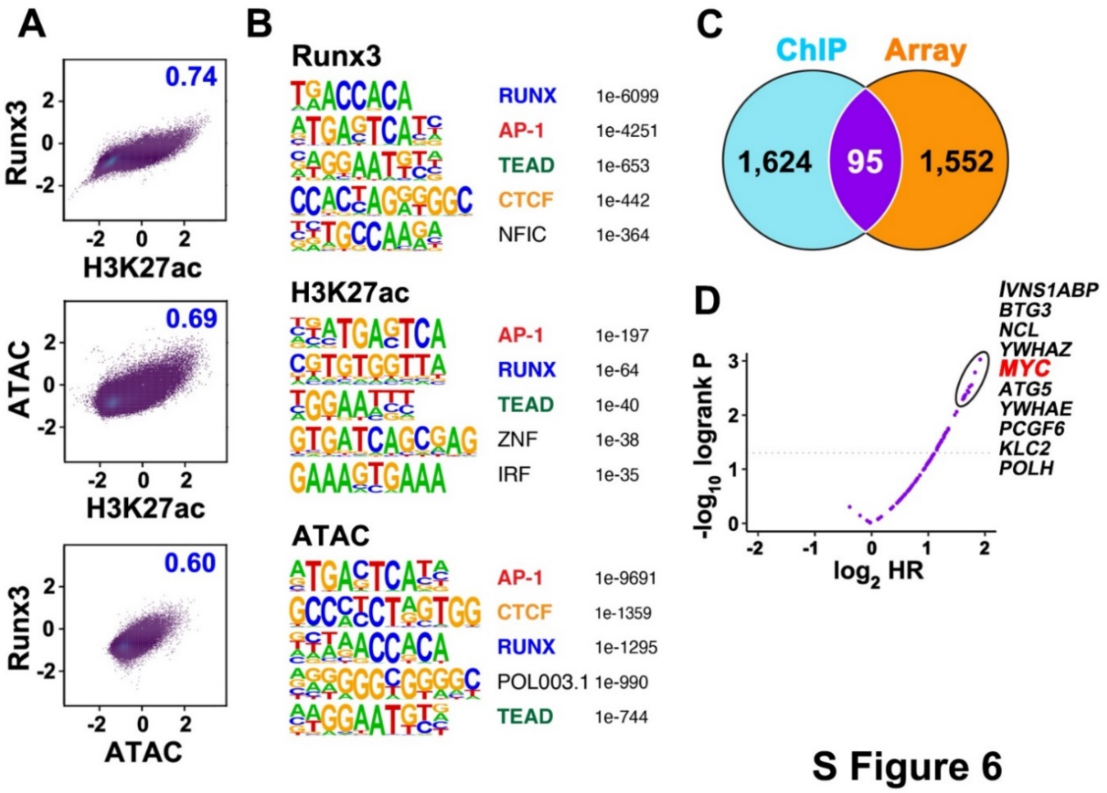


**Fig. S6 Identification of target genes upregulated by Runx3, a transcriptional activator, in *p53*-deficient mOS cells.**

(**A**) Genome-wide correlations among the profiles of Runx3 and H3K27ac ChIP-seq and ATAC-seq (Fig. 2A). Pearson correlation coefficient is shown in each panel. (**B**) Enriched consensus motif of transcription factors in the genome-wide profiles of Runx3 and H3K27ac ChIP-seq and ATAC-seq. (**C**) Number of genes upregulated by Runx3 in BM-MSCs (Array) or bound by Runx3 in mOS cells (ChIP). The 95 overlapping genes, listed in table S2, were considered as Runx3 target genes in this study. (**D**) Volcano plot of the hazard ratio values for the 95 Runx3 target genes, as calculated based on the TARGET data. Myc was among the factors most strongly associated with poor prognosis (circled).

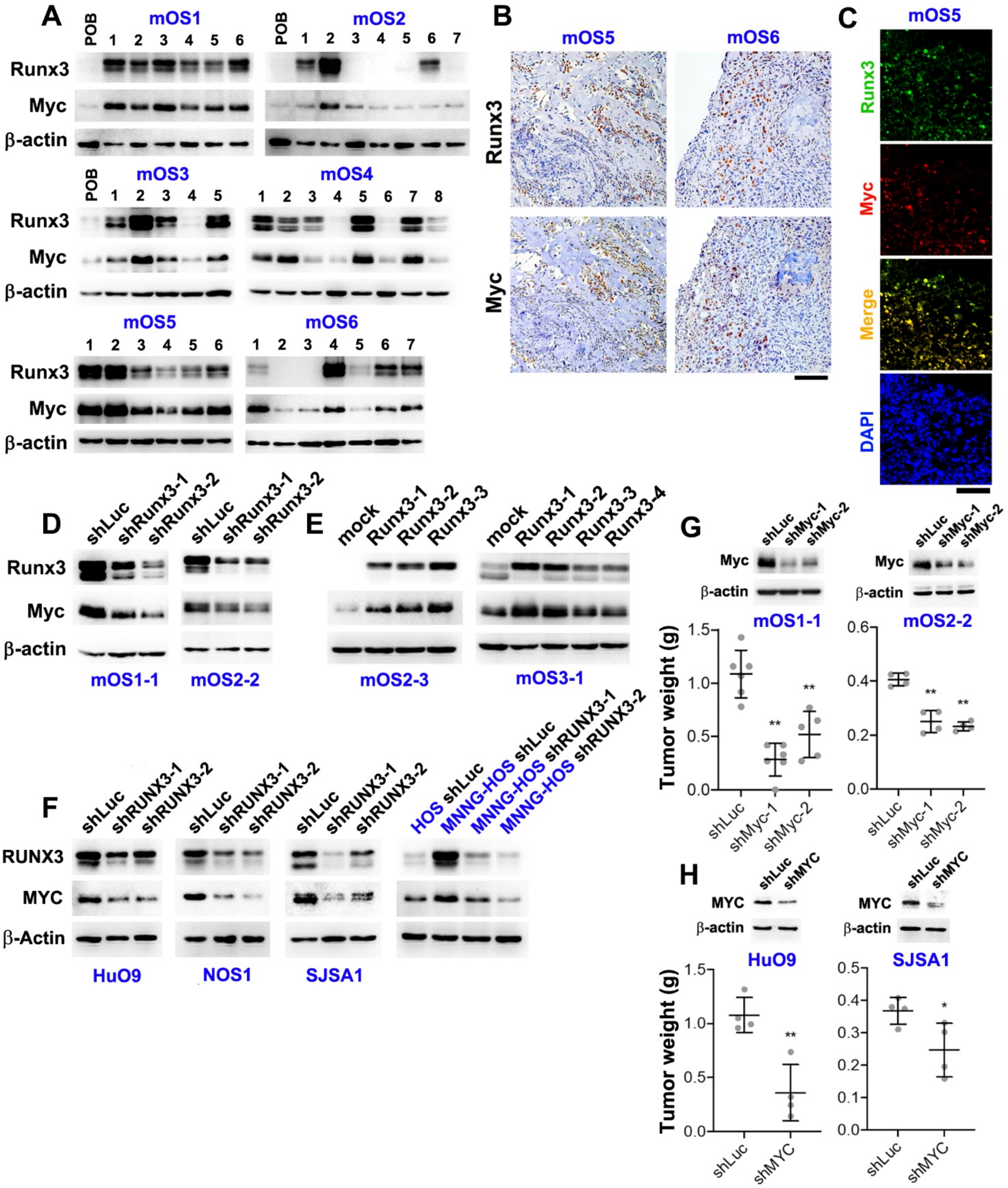
A
mOS1
mOS2
mOS3
mOS4
mOS5
mOS6
Runx3
Myc
β-actin
B
C
D
E
F
G
H
shLuc
shRunx3-1
shRunx3-2
mock
Runx3-1
Runx3-2
Runx3-3
Runx3-4
mOS1-1
mOS2-2
mOS2-3
mOS3-1
shRUNX3-1
shRUNX3-2
RUNX3
MYC
β-Actin
HuO9
NOS1
SJSA1
HOS shLuc
MNNG-HOS shLuc
MNNG-HOS shRUNX3-1
MNNG-HOS shRUNX3-2
Merge
DAPI
shMyc-1
shMyc-2
shMYC
Tumor weight (g)

S Figure 7

**Fig. S7 High correlation between Runx3 and Myc levels in *p53*-deficient OS.**
(**A**) Amounts of Runx3 and Myc in clonal OS cells derived from primary OS (mOS1–6) developed in six individual *OS* mice, as determined by western blotting. Clonal OS cells derived from mOS1–3 were also used in fig. S2C. (**B**) Immunodetection of Runx3 and Myc in mOS5 and mOS6. Upper and lower panels show serial sections. Counterstaining was performed with hematoxylin. Scale bar = 100 µm. (**C**) Co-immunodetection of Runx3 (green) and Myc (red) in mOS5. Nuclei were visualized with DAPI. Scale bar = 50 µm. (**D**) Effect of knockdown of Runx3 on expression of Myc in mOS1-1 (clone 1 of mOS1) and mOS2-2 (clone 2 of mOS2) cells, as determined by western blotting. (**E**) Effects of Runx3 overexpression on Myc expression in mOS2-3 (clone 3 of mOS2) and mOS3-1 (clone 1 of mOS3) cells, as determined by western blotting. (**F**) Effects of knockdown of RUNX3 on MYC expression in human OS cell lines HuO9, NOS1, SJSA1, and MNNG-HOS, as determined by western blotting. (**G**) Effects of knockdown of Myc on tumorigenicity of mOS1-1 and mOS2-2 cells. Levels of the indicated proteins in the cells were determined by western blotting (upper), and tumorigenicity was evaluated by allograft (lower). (**H**) Knockdown effects of MYC on tumorigenicity of HuO9 and SJSA1 cells. Levels of the indicated proteins in the cells were determined by western blotting (upper), and tumorigenicity was evaluated by xenograft (lower). $^{**}p < 0.01$; $^{*}p < 0.05$.

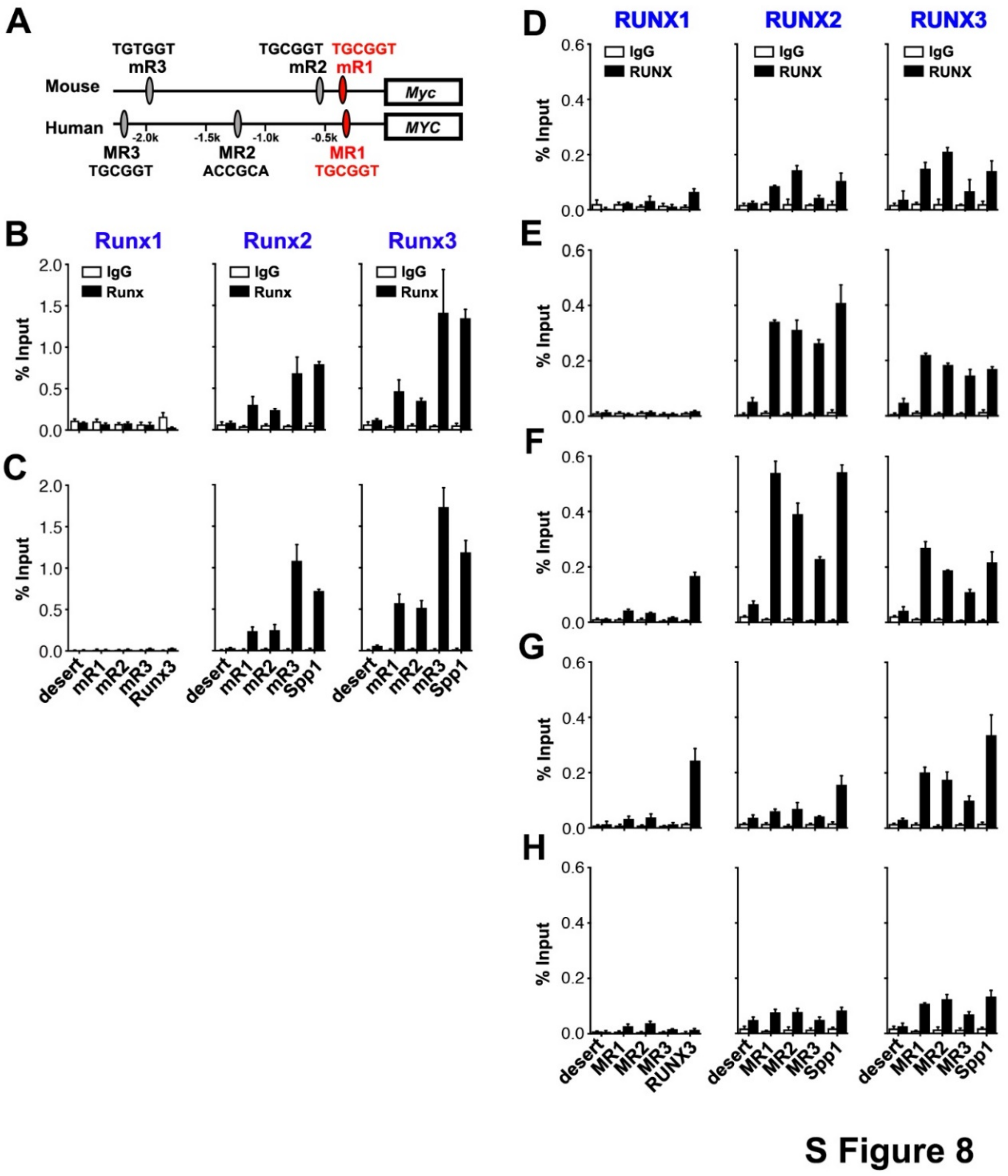


**Fig. S8 Occupancy of Runx proteins in the *Myc/MYC* promoter.**
(**A**) Runx/RUNX consensus binding sites, mR1/MR1, mR2/MR2, and mR3/MR3 in the *Myc/MYC* promoter. (**B** and **C**) Occupancy of Runx1–3 on mR1–3 in mOS1-1 (**B**) and mOS2-2 (**C**) cells, as assessed by ChIP. (**D** to **H**) Occupancy of RUNX1-3 on MR1-3 assessed by ChIP in SJSA1 (**D**), NOS1 (**E**), HuO9 (**F**), MNNG-HOS (**G**), and 143B (**H**) cells. A Runx consensus site in the *Runx3(P1)* promoter (RUNX3) (*60*) and the *Spp1* promoter (Spp1) (*29*) were used as positive controls for Runx1 and Runx2/3 ChIP, respectively, and a gene-desert region of the mouse genome (desert) was used as a negative control for all ChIP assays (**B** to **H**) (n=3).

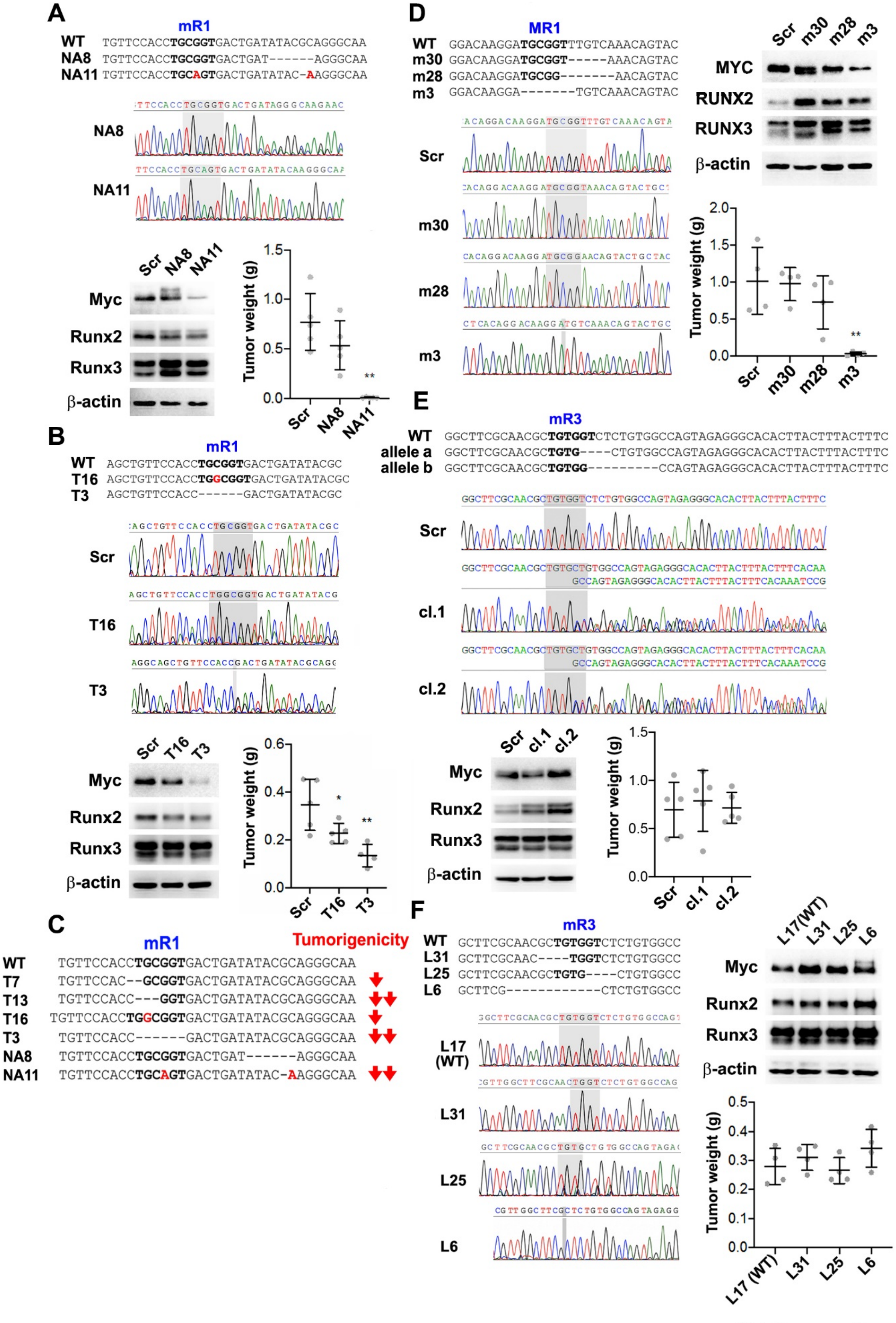
A
mR1
WT TGTTCCACCTGCGGTGACTGATATACGCAGGGCAA
NA8 TGTTCCACCTGCGGTGACTGAT------AGGGCAA
NA11 TGTTCCACCTGCAGTGACTGATATAC-AAGGGCAA
NA8
NA11
Scr NA8 NA11
Myc
Runx2
Runx3
β-actin
Tumor weight (g)
B
mR1
WT AGCTGTTCCACCTGCGGTGACTGATATACGC
T16 AGCTGTTCCACCTGGCGGTGACTGATATACGC
T3 AGCTGTTCCACC------GACTGATATACGC
Scr
T16
T3
Scr T16 T3
Myc
Runx2
Runx3
β-actin
Tumor weight (g)
C
mR1
Tumorigenicity
WT TGTTCCACCTGCGGTGACTGATATACGCAGGGCAA
T7 TGTTCCAC--GCGGTGACTGATATACGCAGGGCAA
T13 TGTTCCACC---GGTGACTGATATACGCAGGGCAA
T16 TGTTCCACCTGGCGGTGACTGATATACGCAGGGCAA
T3 TGTTCCACC------GACTGATATACGCAGGGCAA
NA8 TGTTCCACCTGCGGTGACTGAT------AGGGCAA
NA11 TGTTCCACCTGCAGTGACTGATATAC-AAGGGCAA
D
MR1
WT GGACAAGGATGCGGTTTGTCAAACAGTAC
m30 GGACAAGGATGCGGT-----AAACAGTAC
m28 GGACAAGGATGCGG-------AACAGTAC
m3 GGACAAGGA-------TGTCAAACAGTAC
Scr
m30
m28
m3
Scr m30 m28 m3
MYC
RUNX2
RUNX3
β-actin
Tumor weight (g)
E
mR3
WT GGCTTCGCAACGCTGTGGTCTCTGTGGCCAGTAGAGGGCACACTTACTTTACTTTC
allele a GGCTTCGCAACGCTGTG----CTGTGGCCAGTAGAGGGCACACTTACTTTACTTTC
allele b GGCTTCGCAACGCTGTGG---------CCAGTAGAGGGCACACTTACTTTACTTTC
Scr
cl.1
cl.2
Scr cl.1 cl.2
Myc
Runx2
Runx3
β-actin
Tumor weight (g)
F
mR3
WT GCTTCGCAACGCTGTGGTCTCTGTGGCC
L31 GCTTCGCAAC----TGGTCTCTGTGGCC
L25 GCTTCGCAACGCTGTG----CTGTGGCC
L6 GCTTCG------------CTCTGTGGCC
L17 (WT)
L31
L25
L6
L17(WT) L31 L25 L6
Myc
Runx2
Runx3
β-actin
Tumor weight (g)

S Figure 9

**Fig. S9 mR1 is required for tumorigenicity of *p53*-deficient OS cells.**
(**A**) Sequence alignments of genome-edited mOS1-1 cells, NA8 and NA11, bearing homozygous indels around or within mR1. Levels of the indicated proteins in Scr, NA8, and NA11 mOS1-1 cells were determined by western blotting; tumorigenicity of each cell line is shown. (**B**) Sequence alignments of genome-edited mOS2-2 cells, T16 and T3, bearing homozygous indels in mR1 and the control cells (Scr). Levels of the indicated proteins in Scr, T16, and T3 mOS2-2 cells were determined by western blotting, and the tumorigenicity of each cell line is shown. (**C**) Summary of reduction in tumorigenicity of mOS cells caused by indels around or within mR1 shown in (**A**), (**B**), and Fig. 3, G and J. More or less than 50% reduction in tumorigenicity relative to the control is indicated by two downward arrows or one downward arrow, respectively. (**D**) Sequence alignments of genome-edited SJSA1 cells, m30, m28, and m3, bearing homozygous indels around or within MR1, and control cells (Scr). Levels of the indicated proteins in Scr, m30, m28, and m3 SJSA1 cells were determined by western blotting, and tumorigenicity of each cell line is shown. (**E**) Sequence alignments of genome-edited mOS1-1 cells, clones 1 and 2, both bearing the same pair of alleles (a and b) with indels around or within mR3, and the control cells (Scr). Levels of the indicated proteins in Scr and clones 1 and 2 mOS1-1 cells were determined by western blotting, and tumorigenicity of each cell line is shown. (**F**) Sequence alignments of genome-edited mOS2-2 cells, L31, L25, and L6, bearing homozygous genetic alterations around or within mR3 and the control cells bearing intact mR3 (L17). Levels of the indicated proteins in L17, L31, L25, and L6 mOS2-2 cells were determined by western blotting, and the tumorigenicity of each cell line is shown. $^{**}p < 0.01$; $^{*}p < 0.05$.

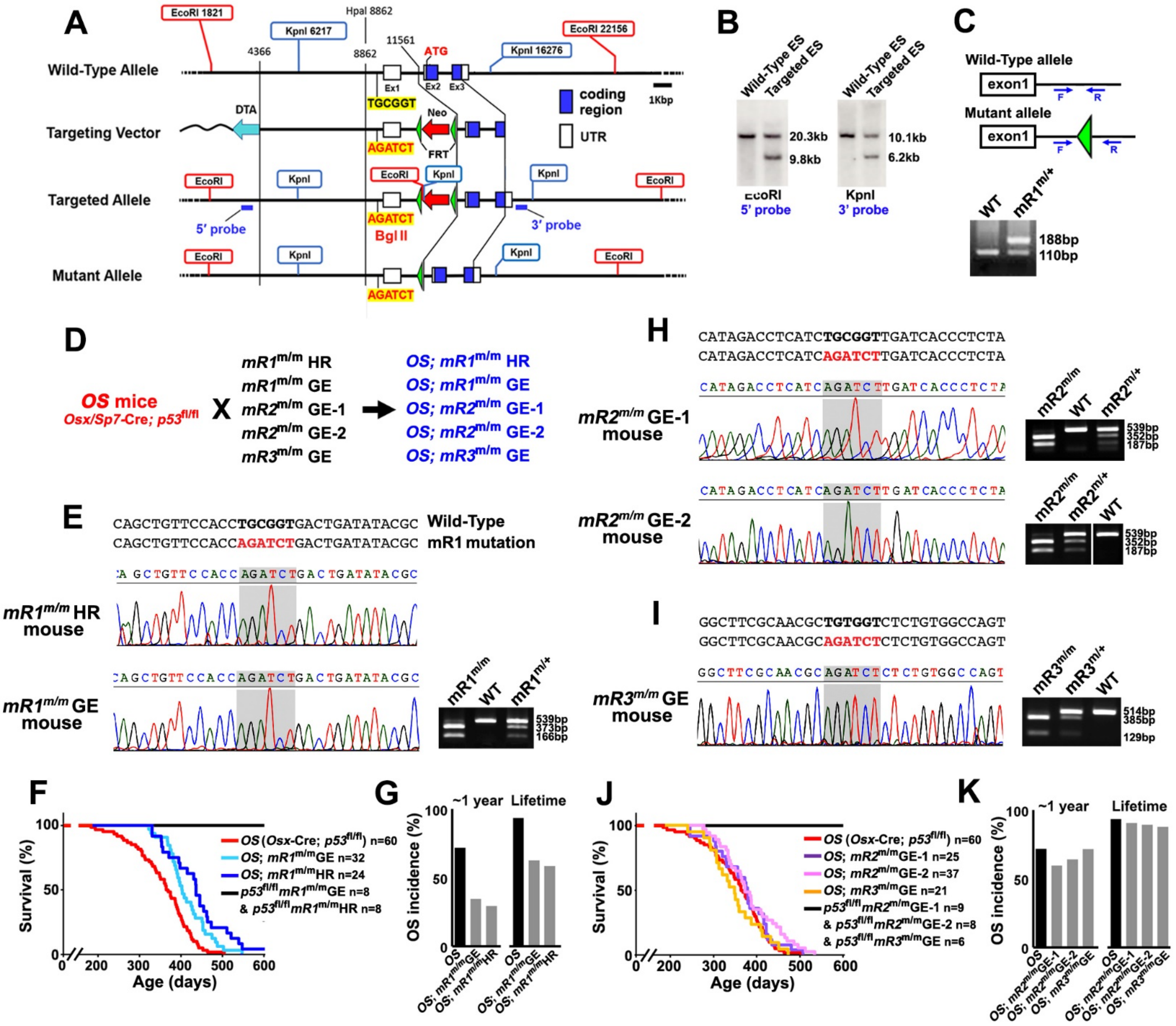
A
Wild-Type Allele
Targeting Vector
Targeted Allele
Mutant Allele
EcoRI 1821
KpnI 6217
HpaI 8862
EcoRI 22156
KpnI 16276
coding region
UTR
1Kbp
DTA
Neo
FRT
5′ probe
3′ probe
Bgl II
B
Wild-Type ES
Targeted ES
20.3kb
9.8kb
10.1kb
6.2kb
EcoRI
5′ probe
KpnI
3′ probe
C
Wild-Type allele
Mutant allele
exon1
188bp
110bp
D
OS mice
Osx/Sp7-Cre; p53fl/fl
E
CAGCTGTTCCACCTGCGGTGACTGATATACGC Wild-Type
CAGCTGTTCCACCAGATCTGACTGATATACGC mR1 mutation
mR1m/m HR mouse
mR1m/m GE mouse
539bp
373bp
166bp
F
Survival (%)
Age (days)
G
~1 year
Lifetime
OS incidence (%)
H
CATAGACCTCATCTGCGGTTGATCACCCTCTA
CATAGACCTCATCAGATCTTGATCACCCTCTA
mR2m/m GE-1 mouse
mR2m/m GE-2 mouse
539bp
352bp
187bp
I
GGCTTCGCAACGCTGTGGTCTCTGTGGCCAGT
GGCTTCGCAACGCAGATCTCTCTGTGGCCAGT
mR3m/m GE mouse
514bp
385bp
129bp
J
Survival (%)
Age (days)
K
~1 year
Lifetime
OS incidence (%)

**S Figure 10**

**Fig. S10 mR1 is required for tumorigenicity of *OS* mice.**

(**A**) Targeting vector for homologous recombination and strategy for generating an mR1-mutated mouse line by *BglII* substitution. (**B**) Southern blot analyses of genomic DNA of wild-type and targeted ES cells digested with *EcoRI* (left) or *KpnI* (right) using the 5' or 3' probes shown in (**A**), respectively. (**C**) Genotyping PCR for detection of wild-type and mutated alleles using F and R primers. (**D**) Generation of *OS; mR1*$^{m/m}$, *OS; mR2*$^{m/m}$, and *OS; mR3*$^{m/m}$ mice. Two independent mouse lines were prepared for each of *mR1*$^{m/m}$ and *mR2*$^{m/m}$. One *mR1*$^{m/m}$ line (*mR1*$^{m/m}$ HR) was generated using homologous recombination, as shown in (**A**) to (**C**), and the other *mR1*$^{m/m}$ line (*mR1*$^{m/m}$ GE), two *mR2*$^{m/m}$ lines (*mR2*$^{m/m}$ GE-1 and -2), and the *mR3*$^{m/m}$ line (*mR3*$^{m/m}$ GE) were generated by genome editing. (**E**) Replacement of mR1 with a *BglII* site in *mR1*$^{m/m}$ HR and GE mice, as confirmed by sequencing. Digestion of 539 bp amplified DNA from the mutant allele with *BglII* produces 373 bp and 166 bp DNA fragments. (**F**) Survival of the indicated mouse lines. *p53*$^{fl/fl}$*mR1*$^{m/m}$ GE and HR lines are Cre-free controls. (**G**) OS incidence in the mouse lines shown in (**F**) within 1 year and throughout the lifespan. The results for *OS* mice are identical to those in Fig. 1, D or E. (**H**) Replacement of mR2 with a *BglII* site in *mR2*$^{m/m}$ GE-1 and -2 mice, as confirmed by sequencing. Digestion of the 539 bp DNA amplified from the mutant allele with *BglII* produces 352 bp and 187 bp DNA fragments. (**I**) Replacement of mR3 with a *BglII* site in *mR3*$^{m/m}$ GE mice, as confirmed by sequencing. Digestion of the 514 bp DNA amplified from the mutant allele with *BglII* produces 385 bp and 129 bp DNA fragments. (**J**) Survival of the indicated mouse lines. *p53*$^{fl/fl}$*mR2*$^{m/m}$ GE-1, -2, and *p53*$^{fl/fl}$*mR3*$^{m/m}$ GE lines are Cre-free controls. (**K**) OS incidence in the mouse lines shown in (**J**) within 1 year and throughout the lifespan. The results of *OS* mice are identical to those in Fig. 1, D or E.

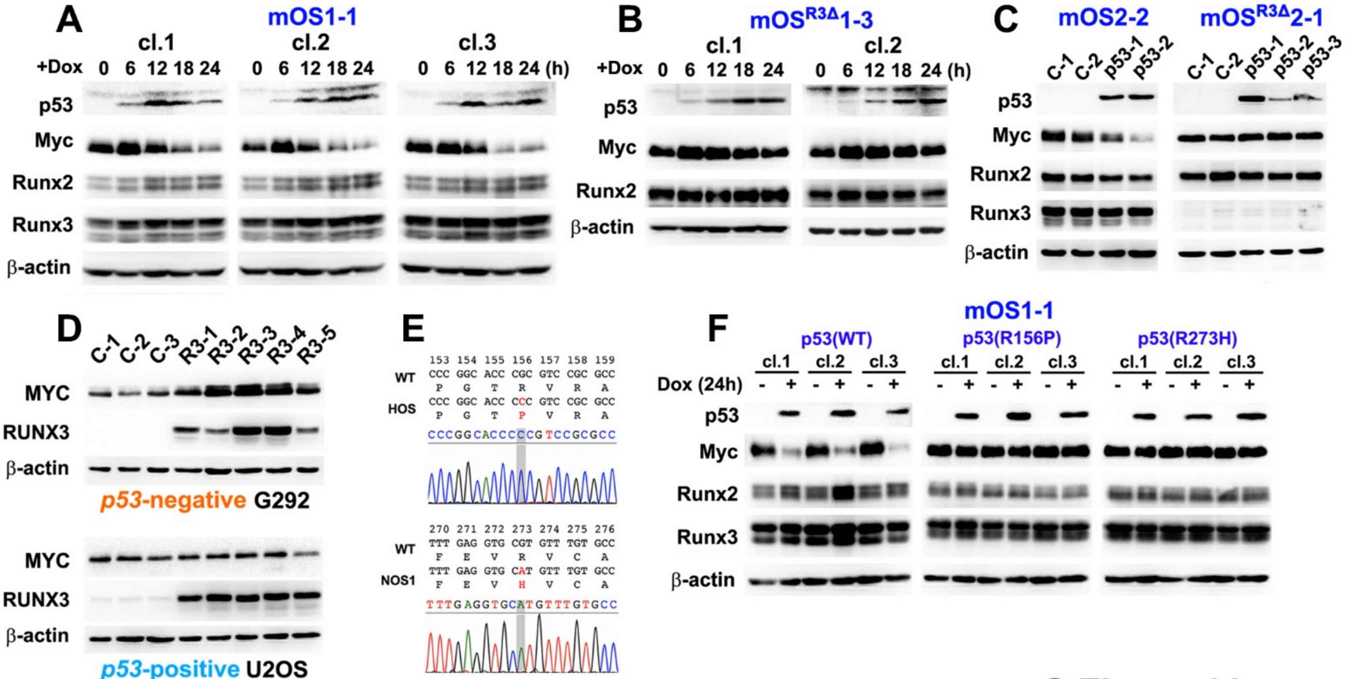


**S Figure 11**

**Fig. S11 p53 attenuates Myc upregulation by Runx3.**

(**A**) Repression of Myc by restoration of p53 induced by doxycycline (Dox) in three clonal mOS1-1 cells (cl. 1–3). Levels of the indicated proteins in the cells were determined by western blotting. (**B**) Myc was not significantly repressed by restoration of p53 induced by Dox in two clonal $mOS^{R3\Delta}1$–3 cells (cl. 1 and 2), which are *Runx3*-negative (fig S13, A and C). Levels of the indicated proteins were determined by western blotting. (**C**) Repression of Myc by restoration of stable p53 in mOS2-2 cells, but not in $mOS^{R3\Delta}2$-1 *Runx3*-negative cells (fig. S13A). (**D**) Levels of the indicated proteins in stable mock (C)- or RUNX3 (R3)-clonal transfectants of *p53*-negative G292 or -positive U2OS cells, as determined by western blotting. (**E**) *TP53* mutations, R156P and R273H, revealed by sequencing of cDNAs expressed in HOS and NOS1 cells, respectively. R156P is inherited in two HOS derivatives, MNNG-HOS and 143B lines, as indicated in the IARC TP53 Database (https://p53.iarc.fr/celllines.aspx). (**F**) Myc was not significantly repressed by the p53(R156P) or p53(R273H) mutants induced by Dox in mOS1-1 cells. Three independent clonal cells (cl. 1–3) were used for each.

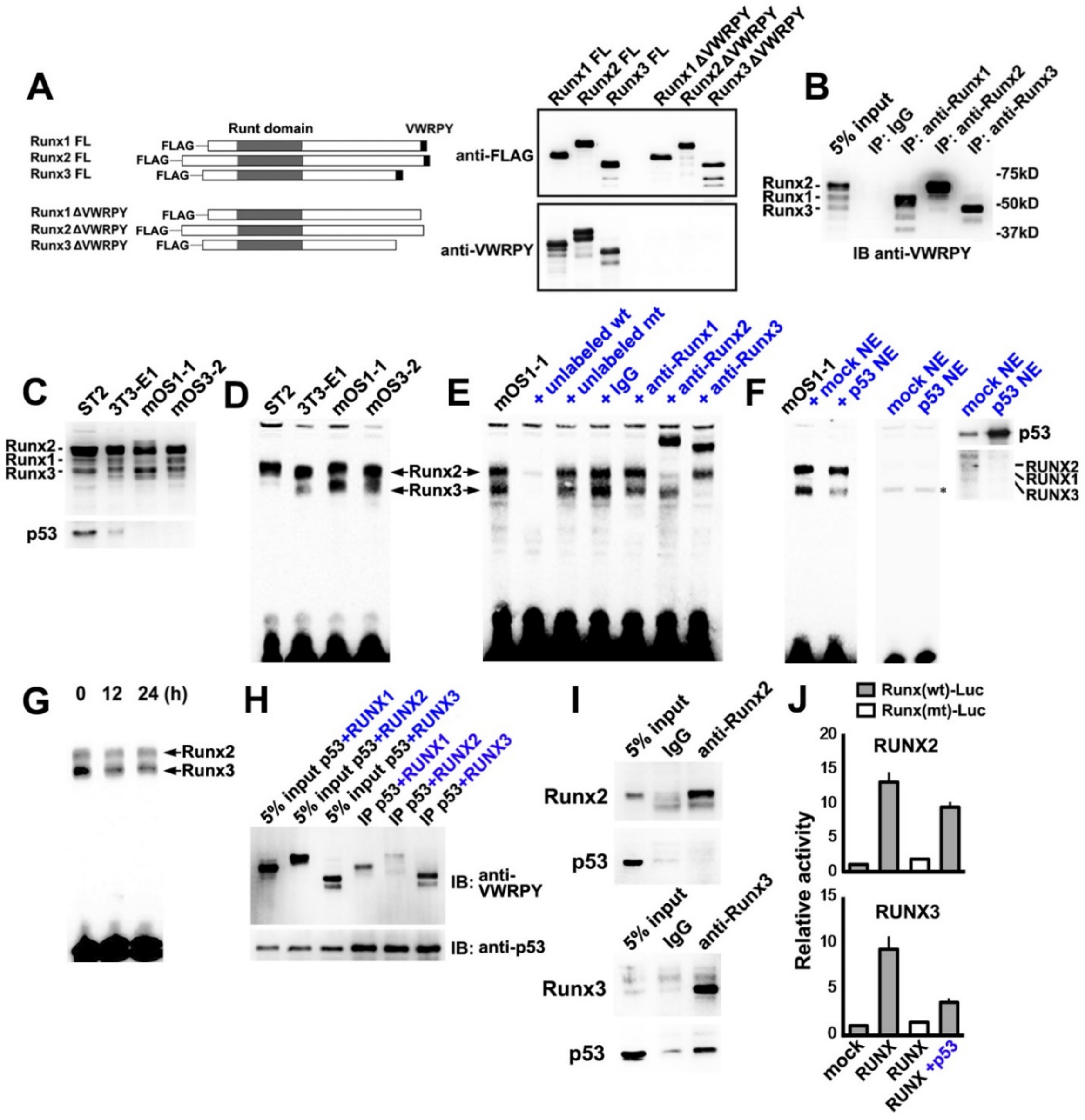
A
Runt domain
VWRPY
Runx1 FL
Runx2 FL
Runx3 FL
FLAG
Runx1 ΔVWRPY
Runx2 ΔVWRPY
Runx3 ΔVWRPY
Runx1 FL
Runx2 FL
Runx3 FL
anti-FLAG
anti-VWRPY
B
5% input
IP: IgG
IP: anti-Runx1
IP: anti-Runx2
IP: anti-Runx3
Runx2
Runx1
Runx3
-75kD
-50kD
-37kD
IB anti-VWRPY
C
ST2
3T3-E1
mOS1-1
mOS3-2
p53
D
E
+ unlabeled wt
+ unlabeled mt
+ IgG
+ anti-Runx1
+ anti-Runx2
+ anti-Runx3
F
+ mock NE
+ p53 NE
mock NE
p53 NE
RUNX2
RUNX1
RUNX3
G
0 12 24 (h)
H
5% input p53+RUNX1
5% input p53+RUNX2
5% input p53+RUNX3
IP p53+RUNX1
IP p53+RUNX2
IP p53+RUNX3
IB: anti-VWRPY
IB: anti-p53
I
IgG
anti-Runx2
anti-Runx3
J
Runx(wt)-Luc
Runx(mt)-Luc
Relative activity
mock
RUNX
RUNX +p53

S Figure 12

**Fig. S12 p53 attenuates DNA binding and transactivation of Runx3.**

(**A**) Pan-reactivity against Runx1–3 proteins of an anti-VWRPY mouse monoclonal antibody established for this study. Exogenously expressed FLAG-tagged RUNX/Runx1–3 (left) was detected specifically and evenly by the anti-VWRPY antibody similarly to the anti-FLAG antibody, as determined by western blotting (right). (**B**) Endogenous amounts and proportions of Runx1–3 in the nuclear extract (NE) of ST2 (a mouse BM-MSC line) cells, detected with the anti-VWRPY antibody (Input). Immunoprecipitates of the NE with specific antibodies against each Runx family member were also subjected to western blotting to identify each Runx in the NE. (**C**) Endogenous amounts of Runx1–3 and p53 proteins in NEs of ST2, 3T3-E1 (a mouse preosteoblastic cell line), mOS1-1, and mOS3-2 (clone 2 of mOS3 shown in fig. S2C). Runx1–3 were detected using the anti-VWRPY antibody. (**D**) EMSA using NEs shown in (**C**) and a labeled DNA probe with an intact mR1 sequence (wt). (**E**) EMSA using NE from mOS1-1 cells, a labeled wt probe, unlabeled wt or mutated mR1 (mt) probes, and normal IgG, anti-Runx1, anti-Runx2, or anti-Runx3 antibodies. The positions of probe-Runx2 and -Runx3 complexes are indicated. (**F**) Inhibition of Runx3–probe binding by p53. NE from mOS1-1 cells was mixed with NE of mock 293T cells or cells expressing exogenous p53, and then subjected to EMSA using a labeled wt probe (left). EMSA using only NE from mock 293T cells or cells expressing exogenous p53 and a labeled wt probe (center). Amounts of exogenous p53 and endogenous RUNX1–3 in NE of 293T cells added to the EMSA were determined by western blotting (right). The asterisk shows a non-specific protein–probe complex. (**G**) EMSA using NE of p53-induced mOS1-1 cl.1 (0, 12, and 24 h) (fig. S11A) and a labeled wt probe. (**H**) Interactions between p53 and RUNX proteins. Immunoprecipitates of whole-cell extracts of 293T cells expressing exogenous p53 and RUNX1, RUNX2, or RUNX3 with an anti-p53 antibody were subjected to western blotting. (**I**) Interactions between endogenous p53 and Runx3 proteins in NE of ST2 cells. Immunoprecipitates of NE of ST2 cells generated with anti-Runx2 or anti-Runx3 antibodies were subjected to western blotting. (**J**) Effect of p53 on transactivation of RUNX2 or RUNX3 in G292 cells (n=3).

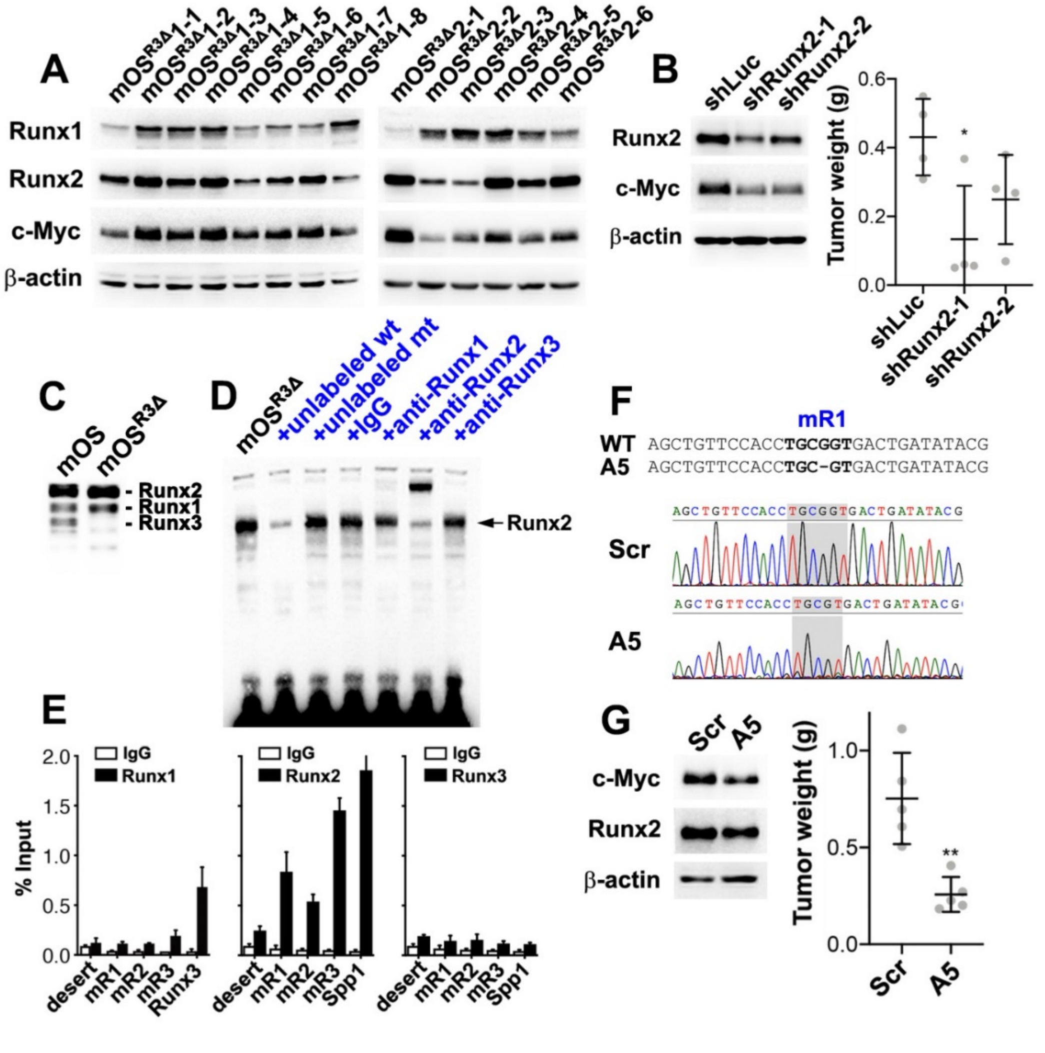


**S Figure 13**

**Fig. S13 Oncogenic role of Runx2 in *Runx3*-deficient mOS cells.**

(**A**) Levels of the indicated proteins in clones of *Runx3*-deficient mOS (mOS$^{R3\Delta}$1 and mOS$^{R3\Delta}$2) cells, which were isolated from OS developed in two individual *OS*; *Runx3*$^{fl/fl}$ mice, as determined by western blotting. (**B**) Effects of knockdown of Runx2 on the tumorigenicity of mOS$^{R3\Delta}$2-1 cells. Levels of the indicated proteins in the cells were determined by western blotting (left), and their tumorigenicity was evaluated by the allograft (right). (**C**) Endogenous levels of Runx1–3 in NE of mOS$^{R3\Delta}$1-3 cells, as determined by western blotting with anti-VWRPY antibody. (**D**) EMSA using NE from mOS$^{R3\Delta}$1–3 cells, labeled wt probe, unlabeled wt or mt probes, and normal IgG, anti-Runx1, anti-Runx2, or anti-Runx3 antibodies. Positions of probe–Runx2 complex is indicated. (**E**) Occupancy of Runx1–3 on each genomic element in mOS$^{R3\Delta}$1–3 cells. (**F**) Genome-edited mOS$^{R3\Delta}$1–3 cell line bearing a 1 bp homozygous deletion in mR1 (A5) and the control line (Scr). (**G**) Levels of the indicated proteins in Scr and A5 mOS$^{R3\Delta}$1-3 cells were determined by western blotting, and the tumorigenicity of each cell line is shown.

**Table S1. Forty-seven common TFs up- or down-regulated in human and mouse OS.**

| | Human OS RNA-seq | | | | | Mouse OS RNA-seq | | | | |
|---|---|---|---|---|---|---|---|---|---|---|
| | Gene | avg. ex. | log2FC | p value | q value | Gene | avg. ex. | log2FC | p value | q value |
| 1 | *BHLHE40* | 12.160 | 1.205 | 0.247 | 0.457 | *Myc* | 11.576 | 1.882 | 0.006 | 0.037 |
| 2 | *JUNB* | 12.110 | 1.518 | 0.113 | 0.276 | *Fos* | 11.397 | 1.103 | 0.141 | 0.388 |
| 3 | *FOS* | 11.765 | 2.508 | 0.248 | 0.458 | *Bhlhe40* | 11.196 | 1.290 | 0.013 | 0.070 |
| 4 | *MYC* | 11.698 | 1.950 | 0.095 | 0.245 | *Etv1* | 10.686 | -1.891 | 0.000 | 0.000 |
| 5 | *LITAF* | 11.357 | 2.025 | 0.004 | 0.023 | *Snai1* | 10.232 | 1.069 | 0.148 | 0.399 |
| 6 | *EBF3* | 10.521 | -1.762 | 0.023 | 0.091 | *Runx3* | 9.962 | 2.166 | 0.001 | 0.009 |
| 7 | *RUNX3* | 9.902 | 4.961 | 0.000 | 0.000 | *Bhlhe41* | 9.806 | 1.070 | 0.139 | 0.385 |
| 8 | *ETV1* | 9.780 | -1.142 | 0.166 | 0.356 | *Litaf* | 9.431 | 1.414 | 0.009 | 0.055 |
| 9 | *IRF7* | 9.709 | 1.135 | 0.300 | 0.515 | *Junb* | 9.416 | 1.370 | 0.015 | 0.080 |
| 10 | *MZF1* | 9.707 | -1.324 | 0.006 | 0.035 | *Heyl* | 9.261 | 1.549 | 0.051 | 0.199 |
| 11 | *BHLHE41* | 9.438 | 2.261 | 0.088 | 0.233 | *Irf7* | 8.879 | 2.574 | 0.000 | 0.004 |
| 12 | *HOXC10* | 9.379 | 2.134 | 0.002 | 0.014 | *Atf3* | 8.810 | 1.647 | 0.091 | 0.291 |
| 13 | *PITX1* | 9.379 | 2.512 | 0.033 | 0.118 | *Prdm1* | 8.582 | 1.096 | 0.185 | 0.454 |
| 14 | *FOSL1* | 9.220 | 1.267 | 0.440 | 0.649 | *Ebf2* | 8.279 | -2.961 | 0.000 | 0.000 |
| 15 | *PRDM1* | 8.655 | 2.610 | 0.004 | 0.024 | *Fosl1* | 8.138 | 3.320 | 0.006 | 0.039 |
| 16 | *HOXC4* | 8.532 | 1.313 | 0.127 | 0.297 | *Nr6a1* | 7.862 | 1.463 | 0.021 | 0.102 |
| 17 | *RFX8* | 8.465 | -1.139 | 0.106 | 0.264 | *Irf8* | 7.852 | 2.690 | 0.000 | 0.001 |
| 18 | *HOXA5* | 8.191 | 1.116 | 0.182 | 0.377 | *Foxd1* | 7.706 | -5.129 | 0.000 | 0.000 |
| 19 | *HOXA7* | 8.159 | 2.510 | 0.007 | 0.036 | *Hoxa3* | 7.484 | 2.516 | 0.002 | 0.016 |
| 20 | *SNAI1* | 8.156 | 4.190 | 0.002 | 0.013 | *Hsf4* | 7.033 | 1.898 | 0.029 | 0.132 |
| 21 | *ATF3* | 8.096 | 2.213 | 0.076 | 0.210 | *Aknad1* | 6.825 | 1.116 | 0.294 | 0.592 |
| 22 | *HOXB3* | 7.892 | 3.696 | 0.000 | 0.002 | *Rorc* | 6.734 | -2.965 | 0.000 | 0.004 |
| 23 | *HOXB2* | 7.885 | 4.429 | 0.000 | 0.000 | *Hoxb2* | 6.672 | 2.544 | 0.022 | 0.106 |
| 24 | *HSF4* | 7.830 | 1.508 | 0.257 | 0.468 | *Pou2f2* | 6.650 | 1.120 | 0.143 | 0.391 |
| 25 | *HOXA3* | 7.783 | 1.324 | 0.246 | 0.455 | *Foxf1* | 5.932 | 1.763 | 0.114 | 0.340 |
| 26 | *EBF2* | 7.770 | -1.064 | 0.109 | 0.270 | *Hoxc10* | 5.895 | 5.879 | 0.000 | 0.002 |
| 27 | *POU2F2* | 7.627 | 4.666 | 0.009 | 0.047 | *Rorb* | 5.890 | -2.016 | 0.038 | 0.161 |
| 28 | *LHX4* | 7.149 | -1.283 | 0.085 | 0.229 | *Hoxa5* | 5.684 | 3.642 | 0.000 | 0.004 |
| 29 | *HEYL* | 7.148 | 8.564 | 0.000 | 0.000 | *Hoxb3* | 5.683 | 2.221 | 0.024 | 0.113 |
| 30 | *SNAI3* | 7.126 | -1.995 | 0.000 | 0.004 | *Hoxc4* | 5.316 | 4.243 | 0.000 | 0.001 |
| 31 | *FOXD1* | 7.044 | -4.595 | 0.000 | 0.000 | *Rfx8* | 5.154 | -1.091 | 0.153 | 0.407 |
| 32 | *HOXC11* | 7.038 | 1.139 | 0.356 | 0.572 | *Hoxa7* | 5.122 | 4.386 | 0.000 | 0.003 |
| 33 | *NR6A1* | 6.999 | 1.537 | 0.079 | 0.218 | *Mzf1* | 5.037 | -1.145 | 0.112 | 0.336 |
| 34 | *RORB* | 6.661 | -1.008 | 0.314 | 0.529 | *Batf2* | 4.894 | 1.634 | 0.050 | 0.196 |
| 35 | *HOXB6* | 6.608 | 4.119 | 0.000 | 0.001 | *Foxp3* | 4.864 | 1.409 | 0.085 | 0.280 |
| 36 | *HOXB7* | 6.540 | 6.186 | 0.000 | 0.000 | *Hoxb4* | 4.829 | 3.468 | 0.001 | 0.007 |
| 37 | *HOXB4* | 6.414 | 4.079 | 0.000 | 0.005 | *Ebf3* | 4.374 | -2.985 | 0.000 | 0.001 |
| 38 | *RORC* | 5.727 | -1.264 | 0.175 | 0.368 | *Pitx1* | 4.230 | 1.268 | 0.624 | 0.856 |
| 39 | *DLX4* | 5.560 | 10.004 | 0.000 | 0.002 | *Hoxb6* | 4.196 | 3.463 | 0.005 | 0.034 |
| 40 | *BATF2* | 5.358 | 4.214 | 0.024 | 0.095 | *Dlx4* | 4.155 | 1.058 | 0.345 | 0.645 |
| 41 | *AKNAD1* | 5.239 | 1.822 | 0.143 | 0.322 | *Hoxc11* | 4.117 | 5.434 | 0.010 | 0.057 |
| 42 | *IRF8* | 5.178 | 6.577 | 0.000 | 0.000 | *Hoxb7* | 3.736 | 4.360 | 0.001 | 0.010 |
| 43 | *SIX6* | 4.414 | -1.825 | 0.140 | 0.318 | *Hoxb5* | 3.318 | 4.126 | 0.001 | 0.009 |
| 44 | *FOXF1* | 4.061 | 5.851 | 0.056 | 0.171 | *Lhx4* | 3.162 | -3.007 | 0.000 | 0.002 |
| 45 | *HOXB5* | 3.941 | 6.620 | 0.000 | 0.001 | *Pou1f1* | 2.437 | -2.329 | 0.008 | 0.048 |
| 46 | *FOXP3* | 3.886 | 2.557 | 0.087 | 0.232 | *Snai3* | 2.402 | -2.155 | 0.027 | 0.123 |
| 47 | *POU1F1* | 2.457 | -1.085 | 0.314 | 0.530 | *Six6* | 0.670 | -1.316 | 0.333 | 0.633 |

**Table S2. Runx3 target genes (95 genes) in the *p53*-deficient context.**

| | | ChIP merged peak | | | | Array |
|---|---|---|---|---|---|---|
| | Gene | Score | Chr | Start | End | log2FC |
| 1 | *Myc* | 314.2 | chr15 | 62328449 | 62328642 | 0.61 |
| 2 | *Gjb4* | 149.8 | chr4 | 127352680 | 127352864 | 1.68 |
| 3 | *Ywhae* | 113.1 | chr11 | 75737056 | 75737217 | 0.39 |
| 4 | *Trmt112* | 81.2 | chr19 | 6909618 | 6909797 | 0.47 |
| 5 | *Got1* | 78.0 | chr19 | 43512028 | 43512212 | 0.44 |
| 6 | *Eef2* | 76.8 | chr10 | 81176443 | 81176577 | 0.39 |
| 7 | *Tm4sf1* | 74.7 | chr3 | 57093681 | 57093865 | 0.53 |
| 8 | *Usp36* | 71.0 | chr11 | 118290235 | 118290448 | 0.81 |
| 9 | *Ube2f* | 70.8 | chr1 | 91250118 | 91250395 | 0.39 |
| 10 | *Eef1a1* | 64.2 | chr9 | 78481855 | 78481973 | 0.30 |
| 11 | *Magohb* | 61.8 | chr6 | 131298615 | 131298766 | 0.55 |
| 12 | *Shoc2* | 59.5 | chr19 | 53944041 | 53944246 | 0.52 |
| 13 | *Ywhaz* | 58.8 | chr15 | 36742686 | 36742978 | 0.63 |
| 14 | *Zdhhc18* | 58.3 | chr4 | 133631581 | 133631805 | 0.58 |
| 15 | *Itpr3* | 56.7 | chr17 | 27056272 | 27056499 | 0.65 |
| 16 | *Cct6a* | 56.6 | chr5 | 129787239 | 129787449 | 0.68 |
| 17 | *Sertad2* | 54.6 | chr11 | 20469468 | 20469705 | 0.60 |
| 18 | *Rps28* | 54.4 | chr17 | 33824438 | 33824659 | 0.38 |
| 19 | *Polh* | 50.3 | chr17 | 46202499 | 46202736 | 0.66 |
| 20 | *Ddx23* | 46.1 | chr15 | 98662903 | 98663030 | 0.39 |
| 21 | *Eif2s3x* | 46.0 | chrX | 94177746 | 94178020 | 0.79 |
| 22 | *Shb* | 44.1 | chr4 | 45473190 | 45473360 | 0.61 |
| 23 | *Ncl* | 44.1 | chr1 | 86359377 | 86359603 | 0.56 |
| 24 | *Fermt2* | 40.8 | chr14 | 45530058 | 45530239 | 0.58 |
| 25 | *Osbpl3* | 40.0 | chr6 | 50355439 | 50355591 | 0.61 |
| 26 | *Ttll7* | 37.7 | chr3 | 147298120 | 147298317 | 0.53 |
| 27 | *Gdnf* | 36.7 | chr15 | 7931711 | 7931903 | 1.34 |
| 28 | *Sla* | 33.9 | chr15 | 66844080 | 66844246 | 1.42 |
| 29 | *Gtf2a2* | 33.9 | chr9 | 70012453 | 70012631 | 0.42 |
| 30 | *Fam49b* | 33.8 | chr15 | 64006172 | 64006373 | 0.65 |
| 31 | *Kbtbd11* | 33.8 | chr8 | 14992834 | 14992999 | 2.15 |
| 32 | *Atg5* | 33.7 | chr10 | 44268231 | 44268490 | 0.32 |
| 33 | *Rpl5* | 33.5 | chr5 | 107938626 | 107938801 | 0.34 |
| 34 | *Cacybp* | 33.2 | chr1 | 160234081 | 160234286 | 0.70 |
| 35 | *Abcf1* | 33.0 | chr17 | 35969721 | 35969911 | 0.43 |
| 36 | *Aebp2* | 32.8 | chr6 | 140780930 | 140781093 | 0.56 |
| 37 | *Lsm5* | 32.6 | chr6 | 56655465 | 56655742 | 0.32 |
| 38 | *Coro1c* | 32.2 | chr5 | 113930979 | 113931142 | 0.66 |
| 39 | *Klhl32* | 31.5 | chr4 | 24788599 | 24788788 | 0.82 |
| 40 | *Tmem161b* | 31.3 | chr13 | 84371156 | 84371323 | 0.41 |
| 41 | *AI314180* | 31.3 | chr4 | 58867381 | 58867490 | 0.31 |
| 42 | *Rrn3* | 31.0 | chr16 | 13780629 | 13780822 | 0.48 |
| 43 | *Senp2* | 30.9 | chr16 | 22076575 | 22076805 | 0.43 |
| 44 | *Zc3h18* | 30.1 | chr8 | 122376531 | 122376696 | 0.39 |
| 45 | *Nras* | 30.1 | chr3 | 103058244 | 103058433 | 0.33 |
| 46 | *Pitpnb* | 30.1 | chr5 | 111228472 | 111228670 | 0.47 |
| 47 | *Loxl4* | 29.3 | chr19 | 42631632 | 42631849 | 1.89 |
| 48 | *Ptk2b* | 29.2 | chr14 | 66221116 | 66221322 | 0.92 |

**Table S2. (continued)**

| | | ChIP merged peak | | | | Array |
|---|---|---|---|---|---|---|
| | Gene | Score | Chr | Start | End | log2FC |
| 49 | *Msn* | 28.6 | chrX | 96086344 | 96086514 | 0.80 |
| 50 | *Ncor2* | 28.6 | chr5 | 125245828 | 125246041 | 0.22 |
| 51 | *Hnrnpa3* | 26.6 | chr2 | 75622336 | 75622482 | 0.43 |
| 52 | *Zfp428* | 26.5 | chr7 | 24507247 | 24507519 | 0.43 |
| 53 | *Abce1* | 26.5 | chr8 | 79679038 | 79679228 | 0.65 |
| 54 | *Hnrnpf* | 25.1 | chr6 | 117895552 | 117895751 | 0.29 |
| 55 | *Zfp706* | 24.2 | chr15 | 36960454 | 36960643 | 0.51 |
| 56 | *Tmem131* | 24.1 | chr1 | 37030761 | 37030932 | 0.37 |
| 57 | *Rpl31* | 22.6 | chr1 | 39349167 | 39349273 | 0.41 |
| 58 | *Fdx1l* | 22.1 | chr9 | 21073490 | 21073676 | 0.43 |
| 59 | *Btg3* | 22.0 | chr16 | 78451490 | 78451682 | 0.32 |
| 60 | *Tbc1d1* | 20.9 | chr5 | 64298395 | 64298615 | 0.32 |
| 61 | *Atp13a3* | 20.8 | chr16 | 30439549 | 30439775 | 0.22 |
| 62 | *Zfp35* | 20.6 | chr18 | 23989537 | 23989641 | 0.41 |
| 63 | *Alkbh8* | 20.6 | chr9 | 3335400 | 3335590 | 0.39 |
| 64 | *Thada* | 20.5 | chr17 | 84466146 | 84466388 | 0.35 |
| 65 | *Snrpc* | 20.1 | chr17 | 27839985 | 27840107 | 0.46 |
| 66 | *Hs6st1* | 18.0 | chr1 | 35935342 | 35935506 | 0.44 |
| 67 | *Gls* | 17.4 | chr1 | 52254535 | 52254792 | 0.79 |
| 68 | *Impad1* | 17.3 | chr4 | 4987635 | 4987754 | 0.39 |
| 69 | *Klc2* | 17.2 | chr19 | 5178643 | 5178837 | 0.77 |
| 70 | *Ivns1abp* | 16.8 | chr1 | 151289579 | 151289733 | 0.45 |
| 71 | *Mif* | 16.7 | chr10 | 75872965 | 75873100 | 0.81 |
| 72 | *Fam72a* | 16.0 | chr1 | 131552046 | 131552165 | 0.47 |
| 73 | *Pvr* | 16.0 | chr7 | 19911334 | 19911544 | 0.92 |
| 74 | *Hnrnpk* | 15.8 | chr13 | 58443938 | 58444049 | 0.27 |
| 75 | *Zmym1* | 15.6 | chr4 | 127061069 | 127061224 | 0.43 |
| 76 | *Rtkn2* | 15.3 | chr10 | 67984319 | 67984548 | 0.81 |
| 77 | *Cdh18* | 15.2 | chr15 | 23682197 | 23682389 | 0.37 |
| 78 | *March6* | 15.1 | chr15 | 31542691 | 31542893 | 0.14 |
| 79 | *Taf5* | 15.0 | chr19 | 47067584 | 47067804 | 0.59 |
| 80 | *Nkrf* | 14.0 | chrX | 36903374 | 36903542 | 0.76 |
| 81 | *Tspan2* | 13.9 | chr3 | 102649230 | 102649367 | 1.34 |
| 82 | *Rpl34* | 13.8 | chr3 | 130827979 | 130828159 | 0.39 |
| 83 | *Fyn* | 12.6 | chr10 | 39335950 | 39336144 | 1.10 |
| 84 | *Siglech* | 12.0 | chr7 | 55539955 | 55540157 | 0.45 |
| 85 | *Rsl24d1* | 11.2 | chr9 | 73165753 | 73165906 | 0.43 |
| 86 | *Pinx1* | 10.3 | chr14 | 63860814 | 63861030 | 0.90 |
| 87 | *Mospd4* | 10.3 | chr18 | 46405082 | 46405294 | 1.33 |
| 88 | *Ecsit* | 9.0 | chr9 | 22090113 | 22090271 | 0.60 |
| 89 | *Panx1* | 8.9 | chr9 | 15045442 | 15045689 | 0.56 |
| 90 | *Anp32b* | 8.8 | chr4 | 46433734 | 46433933 | 0.22 |
| 91 | *Rock2* | 8.5 | chr12 | 16886599 | 16886811 | 0.37 |
| 92 | *Pcgf6* | 7.1 | chr19 | 47050319 | 47050498 | 0.62 |
| 93 | *Ets1* | 6.8 | chr9 | 32629549 | 32629683 | 0.49 |
| 94 | *Ttc14* | 6.7 | chr3 | 33498657 | 33498913 | 0.93 |
| 95 | *Ube2q2* | 5.7 | chr9 | 55126410 | 55126562 | 0.85 |
